# Discovery of homogentisic acid as a precursor in trimethoprim metabolism and natural product biosynthesis

**DOI:** 10.1101/2022.09.22.509022

**Authors:** Andrew C. McAvoy, Paxton H. Threatt, Joseph Kapcia, Neha Garg

## Abstract

Opportunistic infections by *Burkholderia cenocepacia* are life threatening for patients suffering from cystic fibrosis and chronic granulomatous disease. These infections are often associated with variable clinical outcomes, prompting an interest into molecular investigations of phenotypes associated with disease severity. The production of the pyomelanin pigment is one such phenotype, which was recently linked to the ability of clinical strains to carry out biotransformation of the antibiotic trimethoprim. However, this biotransformation product was not identified, and differences in metabolite production associated with pyomelanin pigmentation are poorly understood. Here, we identify several key metabolites produced exclusively by the pyomelanin-producing strains. To provide insight into the structures and biosynthetic origin of these metabolites, we developed a mass spectrometry-based strategy coupling unsupervised *in silico* substructure prediction with stable isotope labeling referred to as MAS-SILAC (Metabolite Annotation assisted by Substructure discovery and Stable Isotope Labeling by Amino acids in Cell culture). This approach led to discovery of homogentisic acid as a precursor for biosynthesis of several natural products and for biotransformation of trimethoprim, representing a previously unknown mechanism of antibiotic tolerance. This work presents application of computational methods for analysis of untargeted metabolomic data to link the chemotype of pathogenic microorganisms with a specific phenotype. The observations made in this study provide insights into the clinical significance of the melanated phenotype.

## Introduction

The bacterial pathogen *Burkholderia cenocepacia* causes fatal infections in immunocompromised humans, particularly in patients suffering from cystic fibrosis (CF) and chronic granulomatous disease.^1–3^ Infections by *B. cenocepacia* are associated with poor clinical outcomes and have the potential to develop into a deadly combination of bacteremia, necrotizing pneumonia, and rapid respiratory failure known as cepacia syndrome.^4^ These pathogens are highly transmissible and are resistant to many standard antibiotic treatments.^5, 6^ Differences in virulence have been noted even between clonally related *B. cenocepacia* strains across multiple infection models, but the molecular basis of these differences is not fully understood.^7, 8^ A subset of *B. cenocepacia* strains produce the pigment pyomelanin due to a G378R amino acid substitution in homogentisate 1,2-dioxygenase (HmgA) within the tyrosine catabolism pathway (Fig. 1A).^9^ In *B. cenocepacia*, pigment production is associated with protection from extracellular oxidative stress *in vitro* and phagocytic killing by macrophages.^10, 11^ We recently showed that pigmented strains can metabolize the antibiotic trimethoprim, increasing the minimal inhibitory concentration of this antibiotic.^12, 13^ Trimethoprim in combination with sulfamethoxazole (co-trimoxazole) is used as one of the first lines of treatment in the clinic.^14^ However, the details of how trimethoprim undergoes metabolic transformation by pigmented strains are not known. Thus, we hypothesized that the metabolic makeup of pigmented strains may be different from their non-pigmented counterparts, and that the characterization of these differences will provide insights into mechanisms underlying antibiotic metabolism as well as virulence in these strains. Comparative statistical analysis of metabolomics data generated in this study supported our hypothesis and enabled selection of metabolite features associated with pigmented strains. However, conventional mass spectral database searching failed to provide annotations for these pigment-specific metabolite features. Here, we utilize recently developed computational methods in mass spectrometry and stable isotopic labeling to demonstrate that sulfur metabolism is inextricably linked to covalent modification of trimethoprim and biosynthesis of several bioactive natural products by pigmented strains through accumulation of homogentisic acid.

**Figure 1.**
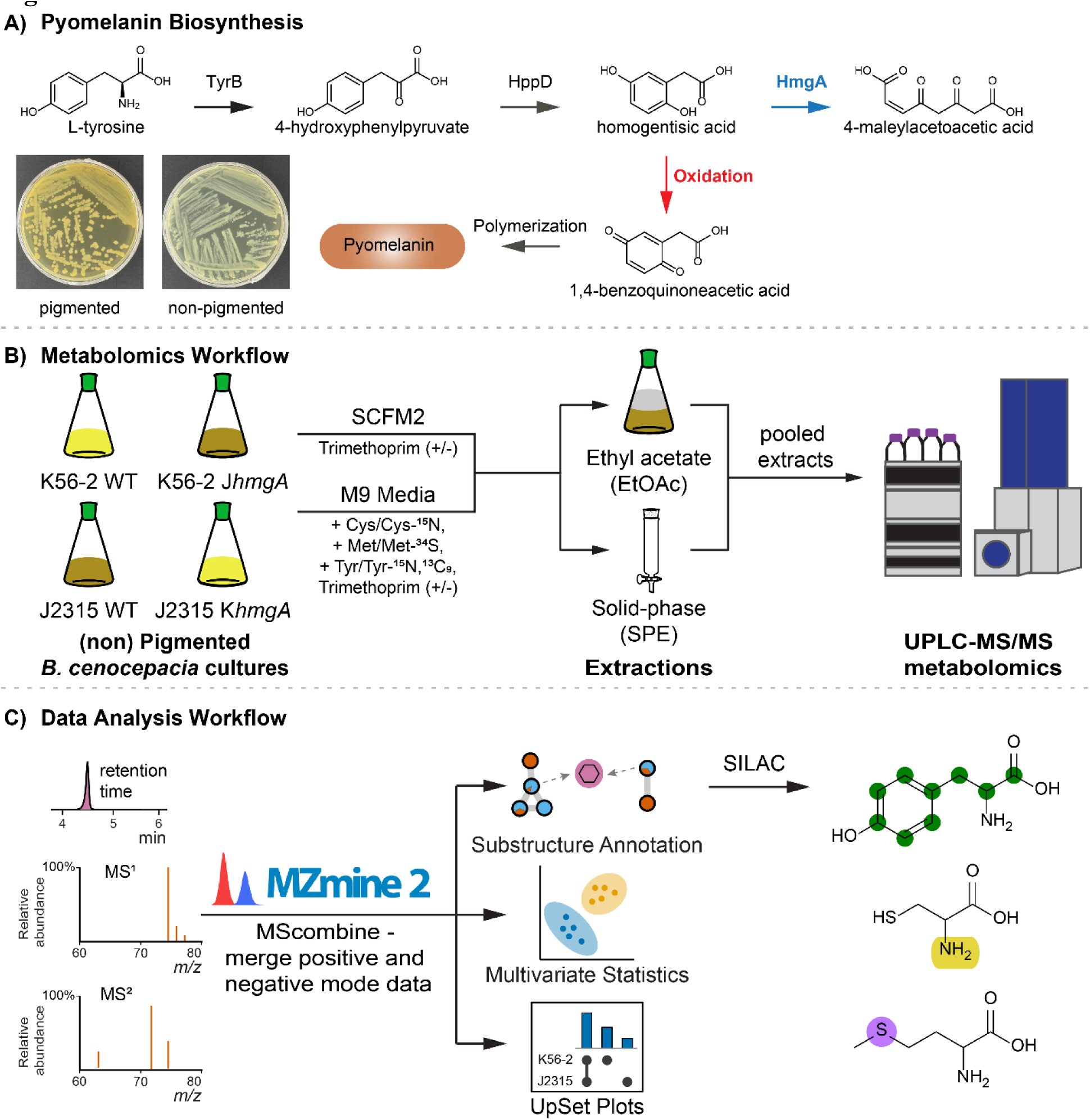
**A**) Biochemical pathway for pyomelanin production in *Burkholderia cenocepacia*. A single amino acid substitution in the gene encoding homogentisic acid 1,2-dioxygenase (HmgA) leads to accumulation of homogentisic acid, which is spontaneously oxidized and polymerized to form pyomelanin in pigmented strains. Overview of **B**) experimental design and **C**) metabolomic data analysis workflow used within the present study.

We refer to our strategy as MAS-SILAC (**<u>M</u>**etabolite **<u>A</u>**nnotation assisted by **<u>S</u>**ubstructure discovery and **<u>S</u>**table **<u>I</u>**sotope **<u>L</u>**abeling by **<u>A</u>**mino acids in **<u>C</u>**ell culture). This approach couples unsupervised *in silico* substructure prediction using mass spectral data with stable isotope labeling. This approach is developed to annotate “known unknowns” (metabolites not annotated in the data being analyzed, but have previously been described in a database or in the literature) and to provide structural insights into true unknown metabolites produced by pigmented *B. cenocepacia*, while also identifying the biosynthetic precursors required for their production. Stable isotope labeling has been long sought for investigations into natural product biosynthesis.^15^ In this study, application of substructure annotation methods combined with prediction of elemental composition using high resolution mass spectrometry allowed us to accurately predict the use of isotopically labeled precursors when no structural information for the detected metabolites was available via spectral library matching. This work reveals a previously unappreciated association between pyomelanin pigmentation and biosynthesis of secondary metabolite production, and describes a mechanism of bacterial biotransformation of trimethoprim by homogentisic acid. This work provides insights into the potential additional functions associated with bacterial pigmentation (which is ubiquitous in nature), and explores the potential of biochemical pathways for pigment production as an important drug discovery target. Furthermore, this study highlights the application of MAS-SILAC in untargeted metabolomic analysis to gain insight into the structures and biological origins of unknown compounds which cannot be annotated through conventional strategies in natural product dereplication.

## Results and Discussion

### Comparative Metabolomics Analyses of Pigmented and Non-pigmented B. cenocepacia

To compare the metabolic makeup of pyomelanin producing (referred to as pigmented) isolates of *B. cenocepacia*, comparative untargeted metabolomics was employed. An abbreviated workflow is shown in Fig. 1B and C. The naturally non-pigmented *B. cenocepacia* K56-2 and pigmented J2315 wild-type (WT) strains were previously isolated from CF patients.^16^ Additionally, genetically modified laboratory strains switched in pigment phenotype were used to filter metabolite features that are strain-specific, but unrelated to pigmentation. These modified strains include pigmented K56-2 and non-pigmented J2315, hereafter referred to as *B. cenocepacia* K56-2 J*hmgA* and *B. cenocepacia* J2315 K*hmgA* (Materials and Methods).^9^ These strains were cultured in synthetic CF sputum medium (SCFM2) with and without trimethoprim in triplicate. Culturing was performed in the presence of antibiotic trimethoprim since it is a known inducer of secondary metabolites produced by *B. cenocepacia* K56-2 and also undergoes biochemical modification by pigmented *B. cenocepacia*, while also being the first line of defense again *B. cenocepacia* infections.^12, 13^ To enhance extraction of chemically diverse metabolites, liquid-liquid and solid-phase extractions were performed. These extracts were pooled together prior to UHPLC-HRMS/MS analysis in positive and negative ionization modes. Metabolite features detected at unique *m/z* and retention times in the raw data were extracted, aligned, and area under the chromatographic peak for each feature was quantified using MZmine2 (Fig. 1C).^17^ The MScombine R package was used to merge positive and negative ion mode data into a single metabolite feature quantification table, simplifying downstream statistical analysis.^18^ Combining the data and removal of features detected in media blanks and solvent controls resulted in a feature table containing 4,656 unique features. The metabolite feature table generated after these pre-processing steps was used to perform data visualization, multivariate statistical analysis, and feature-based molecular networking (FBMN) via the Global Natural Products Social Molecular Networking (GNPS) online platform to compare the metabolomic profiles between pigmented and non-pigmented *B. cenocepacia* strains (described below).^19, 20^

First, we generated an UpSet plot to visualize the distribution of unique features across samples varying by pyomelanin production phenotype and exposure to trimethoprim (Fig. 2).^21^ The majority of features (2,693, or 57.0%) were detected under all conditions. Notably, the UpSet plot revealed that 329 features (7.0%) were exclusively detected in all samples from pigmented strains, 127 (2.7%) were solely observed in pigmented samples cultured with trimethoprim, and 98 (2.1%) features were unique to pigmented samples cultured in the absence of trimethoprim. Altogether, 554 (11.7%) features were exclusively detected in pigmented strains. The UpSet plot analysis highlights that pyomelanin production in *B. cenocepacia* is associated with biosynthesis of unique metabolites exclusive to pyomelanin-producing strains, and that metabolomic profiles are further influenced by the presence of the antibiotic trimethoprim.

**Figure 2.**
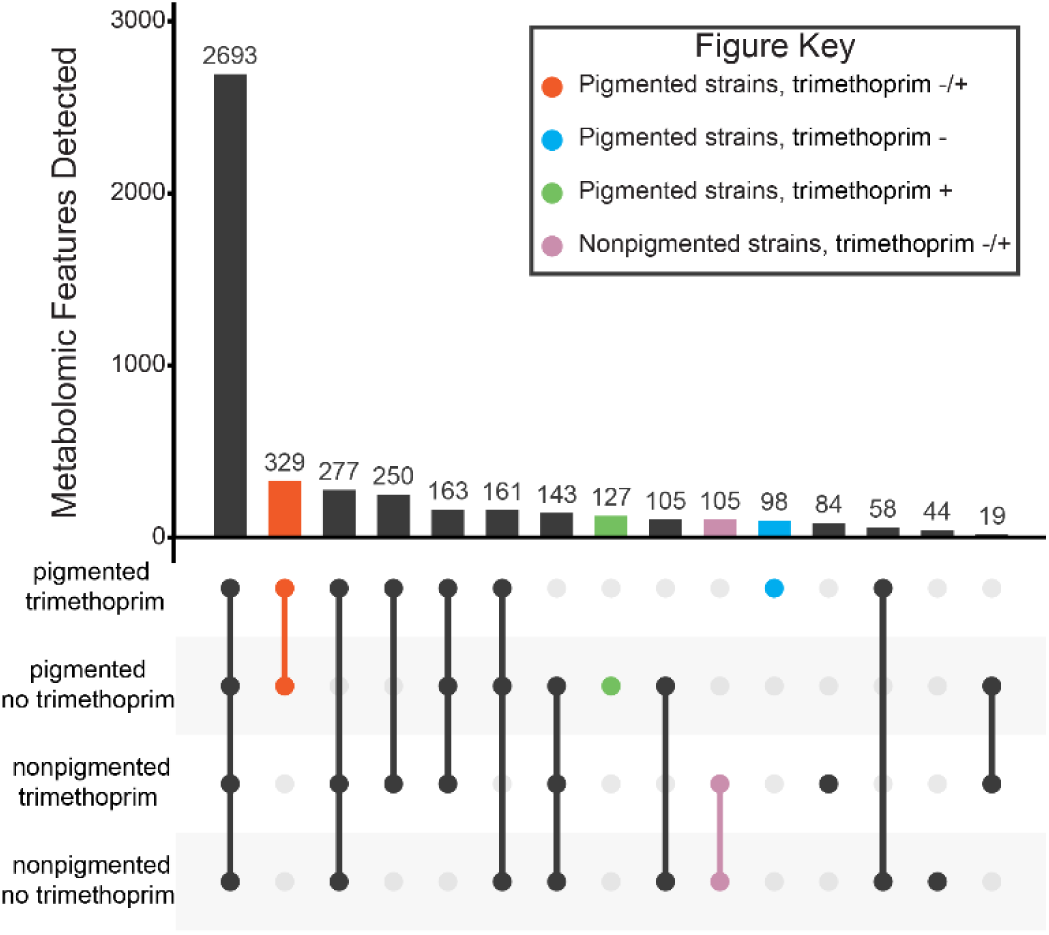
Upset plot displaying the number of features detected in *B. cenocepacia* samples organized by pigment production phenotype and exposure to trimethoprim.

Next, we applied principal component analysis (PCA) as an unsupervised method to visualize whether production of pigment is linked to metabolomic variation. Here, plotting PC1 and PC2 revealed metabolic variation by strain (Fig. 3A), while PC3 revealed variation by pigment phenotype (Fig. 3B). The PC1 and PC2 explained 29.4% of the total variation, with all *B. cenocepacia* J2315 samples (wild-type and non-pigmented mutant) clustering closely together (Fig. 3A). Meanwhile, *B. cenocepacia* K56-2 samples (wild-type and pigmented mutant) formed two additional distinct clusters based on the presence of trimethoprim. Our lab has previously shown that sublethal concentrations of trimethoprim induce a strain-specific response in the K56-2 strain not seen in other *B. cenocepacia* strains, resulting in increased production of secondary metabolites such as fragin and pyrazine-*N*-oxides.^12, 13^ Plotting PC1 against PC3 revealed an additional trend where samples from pigmented strains show separation from those of non-pigmented strains along PC3 (10.9% of the total variance explained) (Fig. 3B). Cross-referencing the PC3 loadings scores with the results of univariate statistical analysis supports this observation (Fig. S1). The features that were detected at significantly higher abundances in pigmented samples have positive PC3 loading scores, while features detected at higher abundances in non-pigmented samples have negative PC3 loadings scores (Fig. S1, Supplementary Data 1). Together, the PCA and UpSet plot indicate that strain-specific metabolites including those produced upon exposure to trimethoprim account for much of the variation in this dataset, while metabolomic differences specific to the pigmented phenotype also contribute to this variation.

**Figure 3.**
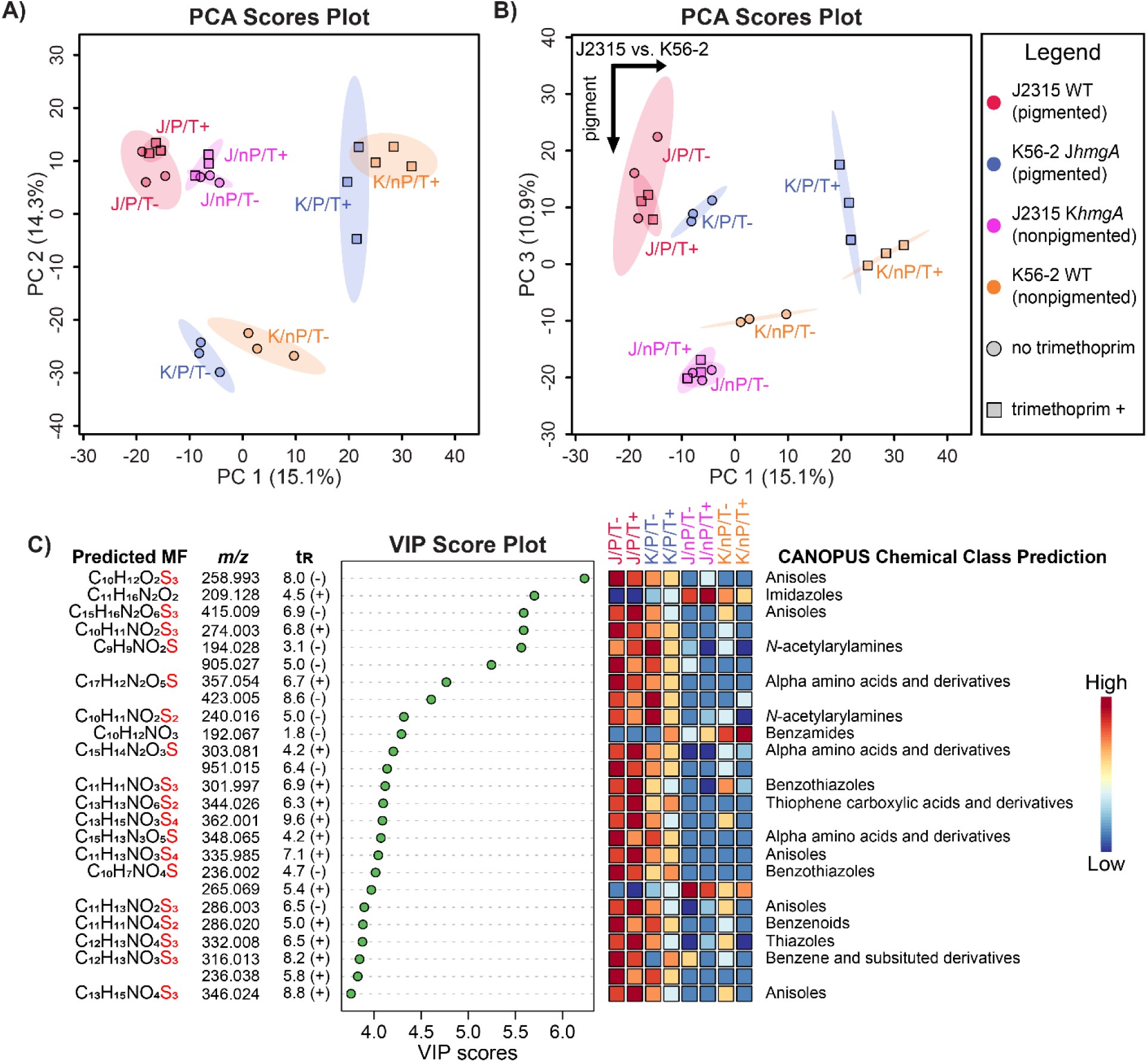
Multivariate analysis of UHPLC-HRMS/MS data collected on pigmented and non-pigmented *B. cenocepacia* strains in the presence or absence of trimethoprim. **A**) PCA scores plot showing variation along PC 1 and PC 2. **B**) PCA scores plot showing variation along PC 1 and PC 3. **C**) VIP score plot displaying the top 25 metabolite features differentially detected across sample groups. VIP features were annotated with the most likely molecular formula (predicted by SIRIUS 4) and ClassyFire compound class as predicted by CANOPUS (when possible). t_R_ refers to retention time and +/− refers to the data acquisition mode (positive or negative) in which the metabolite was detected.

### Annotation of metabolite features specific to pigmented phenotype

We applied partial least-squares discriminant analysis (PLS-DA) as a supervised method to rank metabolites that are differentially detected in strains producing pyomelanin, allowing us to prioritize annotation of phenotype-specific metabolites (Fig. 3C). This method models maximum covariance between variables (i.e. metabolites) and groups (i.e. strains and culture conditions) to best understand factors driving separation. Samples were grouped on the basis of *B. cenocepacia* strain and presence/absence of trimethoprim, resulting in eight distinct groups (Fig. S2). Pigment production and strain-specific differences were clearly associated with larger variation in the PLS-DA model than the differences arising from presence of trimethoprim. A variable importance in projection (VIP) score was calculated for each feature to summarize its contribution to the PLS-DA model (Fig. 3C, Supplementary Data 2). Features with VIP scores greater than one were considered to contribute significantly to the model. These features and the features that show positive contribution to PC3 loading scores in PCA were prioritized for structural annotation. Visualizing the distribution of the highest scoring features across samples cultured under each condition indicated that the majority of top features were exclusively detected in pigmented strains.

Molecular formula predictions by SIRIUS 4 for these features revealed that 18 out of the top 25 features in the score plot contained one or more sulfur atoms and CANOPUS predicted that these compounds contain an aromatic ring (Fig. 3C). All compounds predicted to contain sulfur displayed an [M+2] isotopic peak in the MS^1^ spectra at an intensity consistent with the number of predicted sulfur atoms.

To annotate these metabolites, we first attempted FBMN. In FBMN, each metabolite feature is represented as a node. Structurally-related metabolites are represented as nodes connected by edges. The structural relatedness is established by quantification of similarity between their MS^2^ spectra.^19, 20^ Here, the connected nodes frequently represent chemical transformations of a molecule, enabling propagation of annotations based on *m/z* differences between nodes and manual annotation of MS^2^ peaks. For each feature, consensus experimental MS^2^ spectra are queried against MS^2^ spectral libraries hosted by GNPS to facilitate annotation of known compounds. None of these compounds could be annotated using FBMN or via manual MS^2^ spectral searches across several metabolomic databases. Failure to dereplicate these metabolites motivated the development of MAS-SILAC that combines *in silico* compound annotation with traditional biochemical approaches. The application of this strategy is described below.

MAS-SILAC guides the use of appropriate isotopically labeled amino acids (SILAC) based on the output of unsupervised substructure discovery (metabolite annotation assisted by substructure discovery, MAS) to decode the chemical structures of desired metabolites. The feature with *m/z* 258.993 in negative ion mode (Fig. 3C, corresponding to *m/z* 261.007 in positive ion mode) and predicted molecular formula C_10_H_12_O_2_S_3_ had the highest VIP score in PLS-DA as well as the highest PC3 loadings score (Fig. S1 and Supplemental Data 1). In absence of available annotations, we applied MS2LDA to discover whether a known or unknown substructure is associated with this metabolite.^22^ Using a subset of common fragment peaks and/or neutral losses present in the MS^2^ data, this method discovers chemical substructures referred to as motifs. In the MS2LDA analysis performed on positive mode data, an unknown substructure (motif_584) was discovered for *m/z* 261.007 (Fig. 4A). This unknown substructure motif was present in several additional metabolites across different clusters (Fig. 4B, nodes with blue border). One of the metabolites with *m/z* 551.144 was found to contain a second substructure motif. The chemical structure of this motif was previously annotated as trimethoprim (Fig S3A,B).^12, 13^ Mass spectral annotation of additional nodes in the network revealed that this motif was present in metabolites containing adenine and anthranilic acid as substructures (Figs. S4,S5). Thus, substructure discovery combined with analysis of the molecular network revealed that the nodes in the network correspond to biotransformation products of compounds such as trimethoprim, anthranilic acid, and adenine analogs (Figs. 4,S4,S5, Table S1). Furthermore, the metabolite feature with *m/z* 261.007 is detected at several retention times (Fig. S6). At each of these retention times, a feature with a higher *m/z* containing a product ion with *m/z* 261.007 in the MS^2^ spectrum was also detected. Thus, the metabolite with *m/z* 261.007 is an in-source fragment of a metabolite which is only produced by pigmented strains. In the molecular network in Fig. 4B, the feature with *m/z* 307.013 containing motif_584 was annotated as *m/z* 261.007 with a gain of a carboxylic acid moiety (Fig. S3C). Based on these observations, we hypothesize that the metabolite with *m/z* 307.013 is likely the endogenous metabolite from which the *m/z* 261.007 substructure is derived and serves as the co-substrate involved in biotransformation of compounds such as trimethoprim, adenine, and anthranilic acid. This hypothesis is tested below.

**Figure 4.**
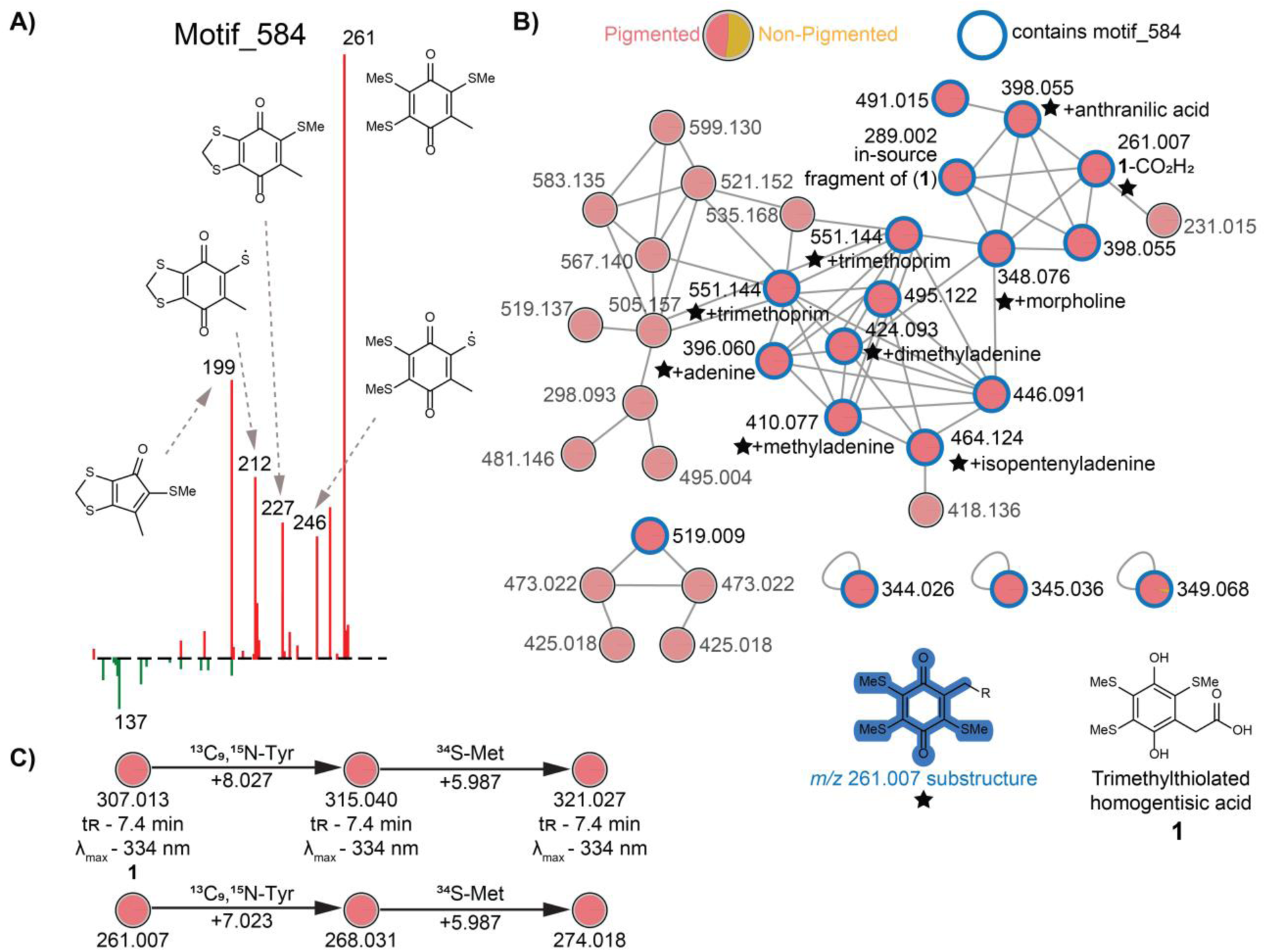
MS2LDA Analysis. **A**) Mass2Motif 584 corresponding to substructure derived from trimethylthiolated homogentisic acid (**1**). **B**) Molecular network representation of features containing motif_584. Pie charts mapped onto nodes represent the relative abundance of the feature across each sample type, as indicated by the integrated area under the chromatographic peak (red: pigmented bacterial samples, gold: non-pigmented bacterial samples). The nodes containing motif_584 are indicated with a blue border. Annotations for biotransformation products are included when available, with stars (★) representing the substructure corresponding to the MS^2^ fragment with *m/z* 261.0072. For the sake of clarity, not all nodes within each cluster are shown. **C**) Representation of observed mass shifts after stable isotope labeling with ^13^C_9_,^15^N-Tyr and ^34^S-Met. Stable isotope labeling was confirmed by matching MS^2^ spectra (Fig. S8), retention time, and UV-visible absorbance spectra.

### Stable Isotope Labeling-Guided Annotation of Biotransformation product of Trimethoprim

Chromatographic analysis of metabolites associated with pigmentation phenotype revealed that the sulfur-containing metabolites, many of which contained the motif 584 and were classified by CANOPUS to contain an aromatic ring, were detected exclusively in the extracts of pigmented *B. cenocepacia* strains. Since pyomelanin production results from a single amino acid substitution in the HmgA enzyme in tyrosine catabolism (Fig. 1A), we reasoned that tyrosine supplementation might be required for their production. Furthermore, given the presence of three sulfur atoms in the chemical formula of the motif 584 substructure (C_10_H_12_O_2_S_3_), we surmised that sulfur-containing amino acids would also be necessary in the biosynthesis. Manual inspection of the MS^2^ spectra from *m/z* 261.007 indicated several fragments separated by neutral losses of 15.024, consistent with the elimination of a methyl radical (Fig. S3C). Such neutral losses are observed during low energy fragmentation of aromatic methoxy- or methylthio-compounds in ESI-MS/MS, which is also why many of these compounds are predicted by CANOPUS as anisoles and benzothiazoles (Fig. 3C).^23–25^ Therefore, guided by the chemical formula corresponding to *m/z* 261.007 (C_10_H_12_O_2_S_3_) as well as predicted chemical classes by CANOPUS, we cultured pigmented strains in minimal media supplemented with tyrosine, methionine, or cysteine. No production of these sulfur-rich metabolites was observed without methionine in M9 minimal medium supplemented with 0.4% glucose, and supplementation with methionine restored production of many of the top metabolite features identified in the VIP plot including *m/z* 261.007 and 307.013 (Fig. S7). Interestingly, these compounds were detected when media was supplemented with methionine alone, but without tyrosine (Fig. S7). Since the shikimate pathway for the biosynthesis of aromatic amino acids is present in *B. cenocepacia*,^26^ we attribute this pathway to serve as a source of tyrosine required for production of compounds in the VIP plot, whereas methionine is the limiting substrate. To confirm that tyrosine and methionine serve as precursors for production of the compounds unique to pigmented strains, we performed a series of stable isotope labeling experiments in which the *B. cenocepacia* strains were cultured in the presence of either labeled or unlabeled tyrosine, methionine, and cysteine (Table 1). The stable isotope labeling data shows that the analyte with *m/z* 261.007 exhibits an increase in mass of 7.024 when cultured in media containing with ^13^C_9_,^15^N-tyrosine and an additional increase of 5.987 in the presence of ^34^S-methionine (Figs. 4C, S8). These shifts in detected *m/z* are consistent with seven carbon atoms originating from tyrosine, and three sulfur atoms derived from methionine. The observed incorporation of labeled carbon atoms from tyrosine indicates that 1,4-benzoquinoneacetic acid provides a carbon backbone for *m/z* 261.007 (Figs. 4C, S8, bottom panel), which was also supported by shifts in specific MS^2^ fragments. Additionally, the observed incorporation of sulfur from isotopically labeled methionine implies that *m/z* 261.007 contains three methylthio-groups. A strong foul-smelling odor was noted in cultures containing methionine, suggesting that methionine is catabolized to methanethiol *in situ*. Methanethiol has been proposed to undergo Michael addition to compounds containing a quinone moiety to yield methylthio-derivatives.^27, 28^ Headspace GC-MS analysis confirmed that dimethyl disulfide (known to be spontaneously formed by oxidative dimerization of methanethiol) is detected in *B. cenocepacia* cultures grown in the presence of methionine (Fig. S9).^29, 30^ To provide further support for methanethiol as the source of methylthio-groups, we cultured the pigmented *B. cenocepacia* J2315 wild-type strain in M9 minimal medium supplemented with ethionine instead of methionine. UHPLC-HRMS/MS analysis of the resulting extracts revealed production of ethylthiolated analogs of pigment specific methylthio-containing compounds (Fig. S10). L-methioninases catalyzing the α, γ-elimination of L-methionine to methanethiol, 2-oxobutanoate, and ammonia are widespread across many bacterial species.^31^ Performing a BLASTP search of L-methioninases (also known as L-methionine γ-lyase) from *Pseudomonas putida* against the genomes of *B. cenocepacia* J2315 and K56-2 yielded five candidate matches (Table S2). These include two putative cystathionine β-lyases, which have also been demonstrated to convert methionine to methanethiol.^32^ In light of this biosynthetic and MS-based evidence supported by isotope labeling, we propose annotating *m/z* 307.013 as trimethylthiolated homogentisic acid, which is derived from tyrosine and methionine (**1**, 3MT-HA). Thus, the fragment with *m/z* 261.007 is trimethylthiolated toluquinone (Fig 4A), where toluquinone has been shown to be produced by the decarboxylation of homogentisic acid.^33^

These observations enable the proposition of the following biosynthetic scheme for production of **1** and inactivation of trimethoprim by *B. cenocepacia* (Fig. 5). Here, the G378R mutation in the HmgA enzyme leads to accumulation of homogentisic acid, which is oxidized to 1,4-benzoquinone.^9^ Next, methanethiolate (derived from thiol-containing compounds such as methionine) undergoes nucleophilic addition to the benzoquinone, as has been proposed to occur *en route* the biosynthesis of methylthio-containing quinone natural products 19-S-methylgeldanamycin and 17,19-dimethylthioherbimycin A.^27, 28^ The resulting intermediate aromatizes and undergoes two additional cycles of oxidation/methanethiolate addition to yield **1**. The antibiotic trimethoprim is known to undergo bioactivation both *in vivo* and *in vitro*.^34^ The activated trimethoprim iminioquinone methide intermediate has been characterized *in vitro* via entrapment with *N*-acetyl cysteine. Addition of *N*-acetyl cysteine takes place at either the C7 *exo*-cyclic methylene group or the C6 carbon on the pyrimidine ring resulting in inactivation of trimethoprim.^34^ Based on this precedence in literature, we propose addition of trimethylthiolated homogentisic acid to bioactivated trimethoprim at these positions resulting in chemical transformation of trimethoprim. Upon culturing in the presence of labeled tyrosine (the precursor of homogentisic acid), expected mass shifts in the biotransformation products of trimethoprim were observed supporting this proposal (Figs. 5B, S11).

**Figure 5.**
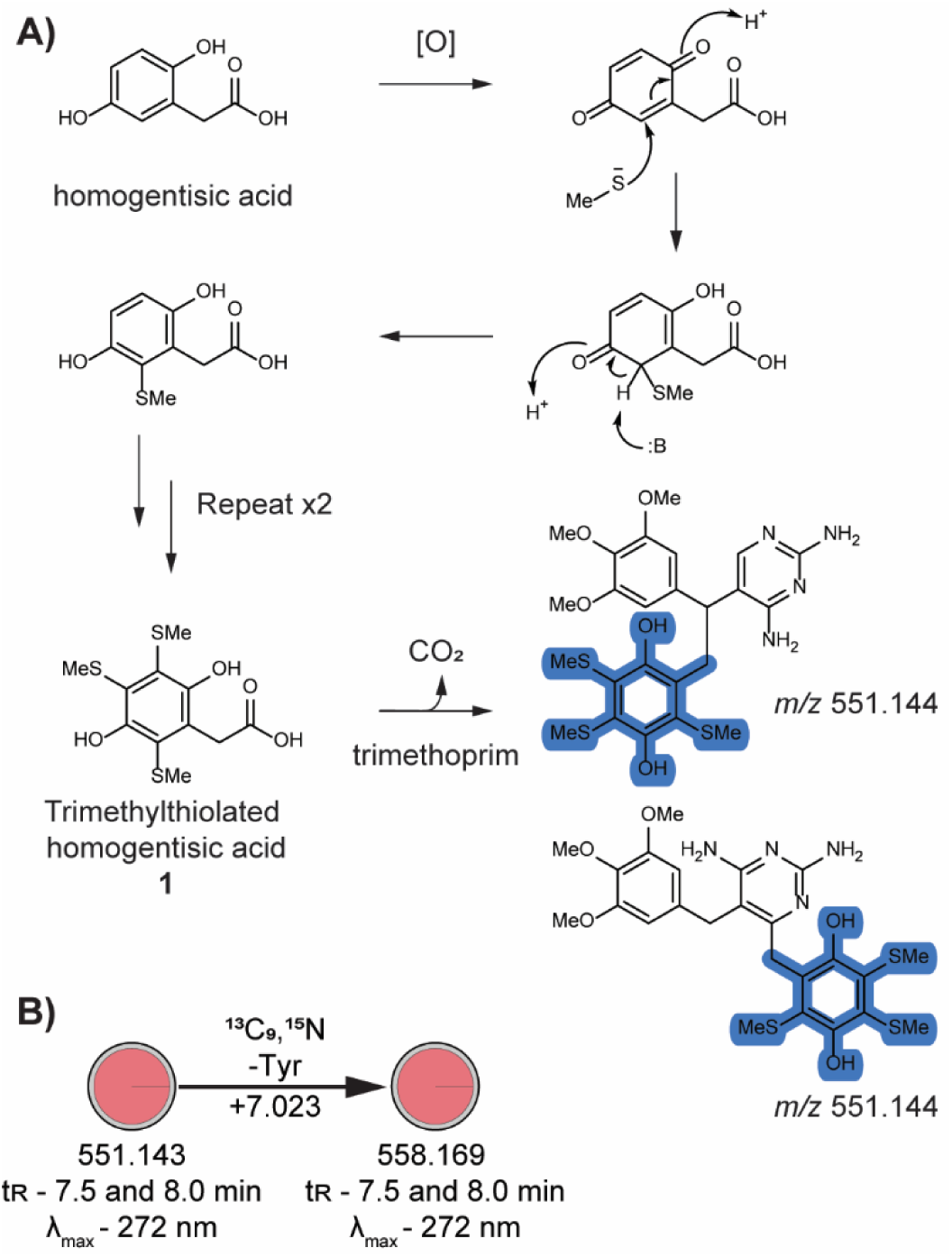
**A**) Proposed biosynthesis of **1** and trimethoprim biotransformation product with *m/z* 551.144. **B**) Representation of observed mass shifts after stable isotope labeling with ^13^C_9_,^15^N-Tyr. Stable isotope labeling was confirmed by matching MS^2^ spectra (Fig. S11), retention time, and UV-visible absorbance spectra.

**Table 1.**
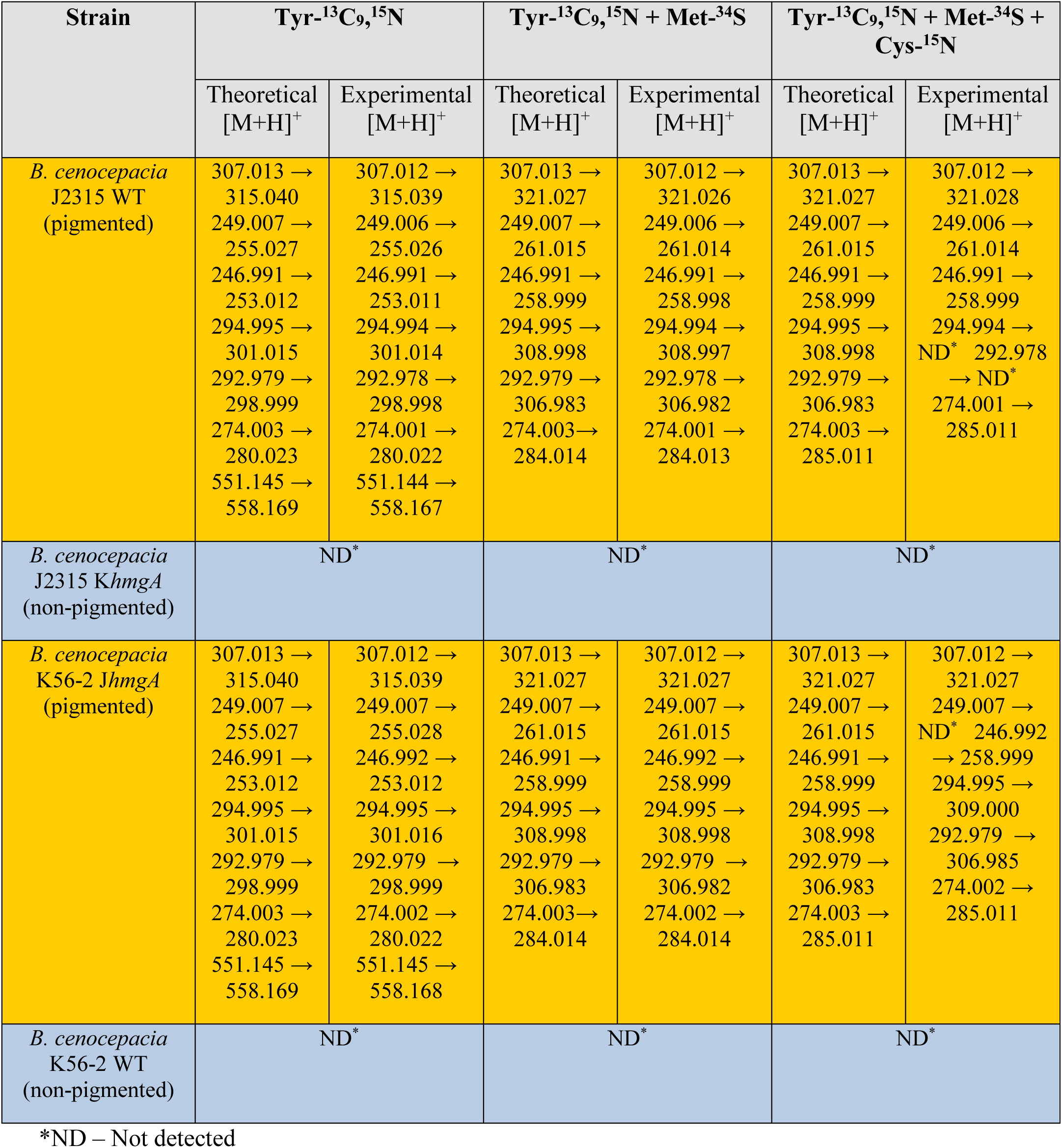
Observed mass shifts for selected pigment-specific metabolites during isotope labeling experiments.

### Synthesis of Trimethylthiolated Homogentisic Acid and Biotransformation Product of Trimethoprim

To further verify that methanethiol and homogentisic acid can result in production of *m/z* 307.013, we attempted to synthesize this compound using methanethiol and homogentisic acid. Although a complex mixture was obtained, **1** was synthesized by addition of methanethiolate to homogentisic acid as one of the products (Fig. 6). The identity of **1** was confirmed by matching retention time, precursor mass, and MS^2^ spectra with the metabolite feature detected in the bacterial extracts (Fig S12A). Additionally, the bacterial metabolite exhibited an absorption signal with λ_max_ at 334 nm that was also observed for the synthetic **1** product. We next attempted synthesis of the biotransformation product at *m/z* 551.145 by first oxidizing trimethoprim to a reactive iminoquinone methide intermediate through addition of sodium hypochlorite as previously described.^34^ Next, activated trimethoprim solution was combined with the reaction mixture described above for the synthesis of **1**. Formation of *m/z* 551.145 was confirmed by matching retention time, precursor mass, and MS^2^ spectra as well as UV-visible absorption spectra with the metabolite feature detected in the bacterial extracts (Fig S12B). The bacterial biotransformation product featured an absorption signal with λ_max_ at 272 nm that was also observed in the synthetic product. Due to formation of a complex mixture of products, the putative chemical structures proposed for **1** and biotransformation product at *m/z* 551.145 were not supported by NMR.

**Figure 6.**
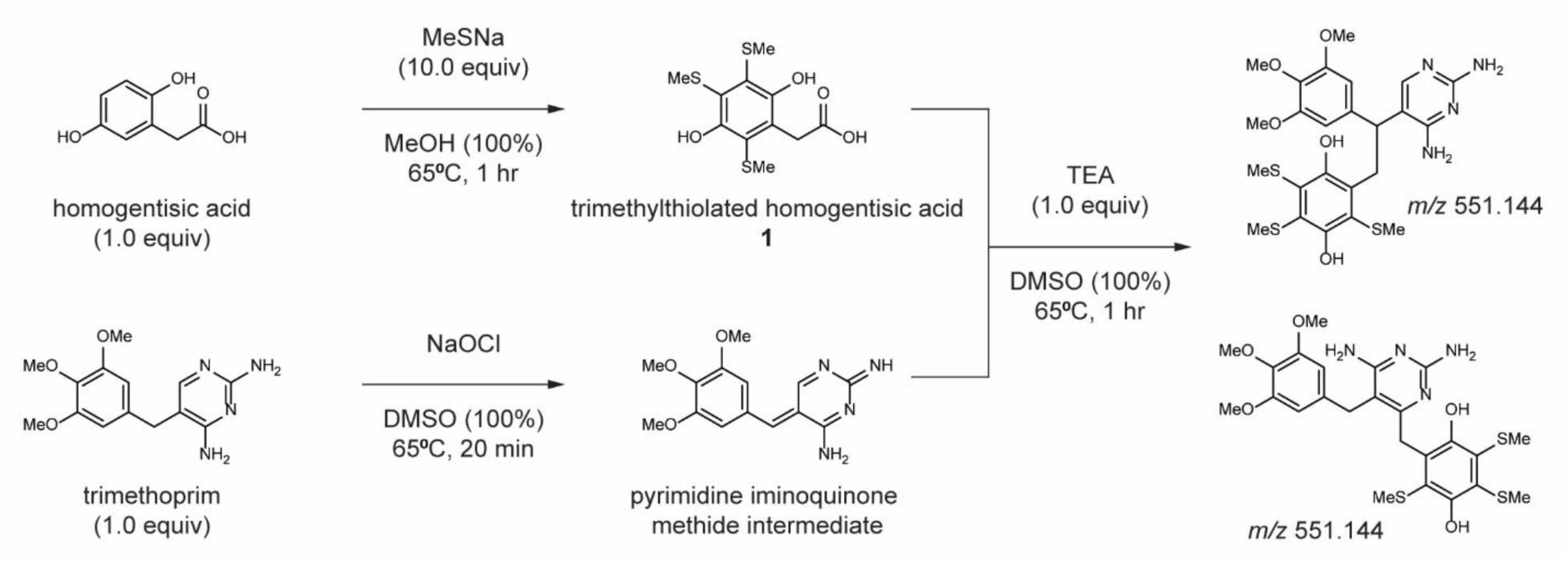
Synthetic Reaction Conditions for 3MT-HA (**1**). The MS^2^ spectral, retention time, and UV-visible absorbance spectral comparison of bacterially produced compounds and the synthetic products are shown in Fig. S12.

Many pathogenic strains of microbial genera produce pyomelanin including *Pseudomonas*, *Ralstonia*, *Legionella*, *Shewanella*, *Serratia*, *Klebsiella, Alcaligenes*, *Enterobacter*, *Hyphomonas*, *Ralstonia*, *Vibrio* and *Acinetobacter* as well as fungi such as *Aspergillus* and *Yarrowia*.^35, 36^

Genetic diversification during infection and harsh conditions such as increased exposure to UV radiation and heavy metals also results in hyperpigmentation.^37^ Thus, it is tempting to propose that homogentisic acid derived biotransformation of trimethoprim may be one of the mechanisms for inactivation of trimethoprim among pyomelanin producing bacteria, therefore meriting further study. A transposon library screening approach will aid in identification of enzymes involved in this biotransformation, which will serve as additional drug targets. We had previously linked the biotransformation of trimethoprim to an increase in the minimal inhibitory concentration of this antibiotic against clinical *B. cenocepacia* strains.^12, 13^ The direct link between pyomelanin biosynthesis and biochemical modification of the antibiotic trimethoprim suggests that inhibition of pyomelanin production in conjunction with antibiotic treatment can also be investigated as a treatment strategy for infections by pyomelanin-producing *B. cenocepacia*. For example, 2-[2-nitro-4-(trifluoromethyl) benzoyl]-1,3-cyclohexanedione (NTBC) is an FDA-approved drug which inhibits the 4-hydroxyphenylpyruvate dioxygenase (Hppd, Figure 1) enzyme responsible for producing homogentisic acid.^38^ Indeed, NTBC has been previously explored as treatment strategy to enhance sensitivity of pyomelanin producing pathogens to oxidative stress.^38^ It stands to reason that inhibition of Hppd by NTBC could increase susceptibility of pigmented *B. cenocepacia* to trimethoprim and block any other potential fitness advantage conferred by pigment-specific metabolites. However, additional studies using animal models would be needed to determine if NTBC administration does result in improved clinical outcomes.

### Stable Isotope Labeling-Guided Annotation of Metabolite Features Specific to Pigmented Strains

The substructure of metabolite with *m/z* 246.992 was annotated as a “loss of methyl group – indicative for presence of methoxy [O-CH_3_] methylsulfyl [S-CH_3_] or methyl group connected to aromatic ring system”. This observation along with the prediction of elemental formula as C_9_H_10_O_2_S_3_ suggested that this compound may also originate from **1**. Additionally, another feature exclusively produced by pyomelanin-producing strains (*m/z* 292.979, t_R_ = 10.6 min, predicted molecular formula: C_10_H_12_O_2_S_4_) was identified as a potential hit to 2,3,5,6-tetrakis(methylsulfanyl)cyclohexa-2,5-diene-1,4-dione (referred to as T1801 D in literature) in the output of *in silico* tool SIRIUS 4 with CSI:FingerID.^39^ T1801 D is part of a family of tri- and tetra-(methylthio) derivatives of hydroquinone (T1801 A and C, **2** & **4**) and *p*-benzoquinone (T1801 B and D, **3** & **5**) previously discovered by Kumagai *et. al* to be produced by a bacterial strain identified as *Pseudomonas* sp. SC-1801.^39^ **2**–**5** exhibit broad antimicrobial activity against Gram-positive bacteria as well as certain fungi.^39^ By searching for features based on *m/z*, candidate features were found for all four compounds within the dataset. All four features were detected exclusively in the pigmented strains (Fig. 7). The MS^2^ fragmentation patterns for these features support the putative annotations of **2**–**5** (Figs. S13-16). Furthermore, these features exhibit distinguishing UV absorption signals with λ_max_ at 333 (**2**), 377 (**3**), 348 (**4**), and 393 nm (**5**), as previously reported for **2**–**5**.^39^ Stable isotope labeling studies also indicated that these metabolites display the expected mass shift under each media condition (Figs 7, S13–16). Consequently, we annotated *m/z* 249.007 as T1801 A (**2**), *m/z* 246.992 as T1801 B (**3**), *m/z* 294.995 as T1801 C (**4**), and *m/z* 292.979 as T1801 D (**5**). To our knowledge, the current study is the first time these antimicrobial natural products have been detected in *Burkholderia* spp. Moreover, this is also the first discussion of the biosynthetic origin of these secondary metabolites and their production in relation to the presence of pyomelanin.

**Figure 7.**
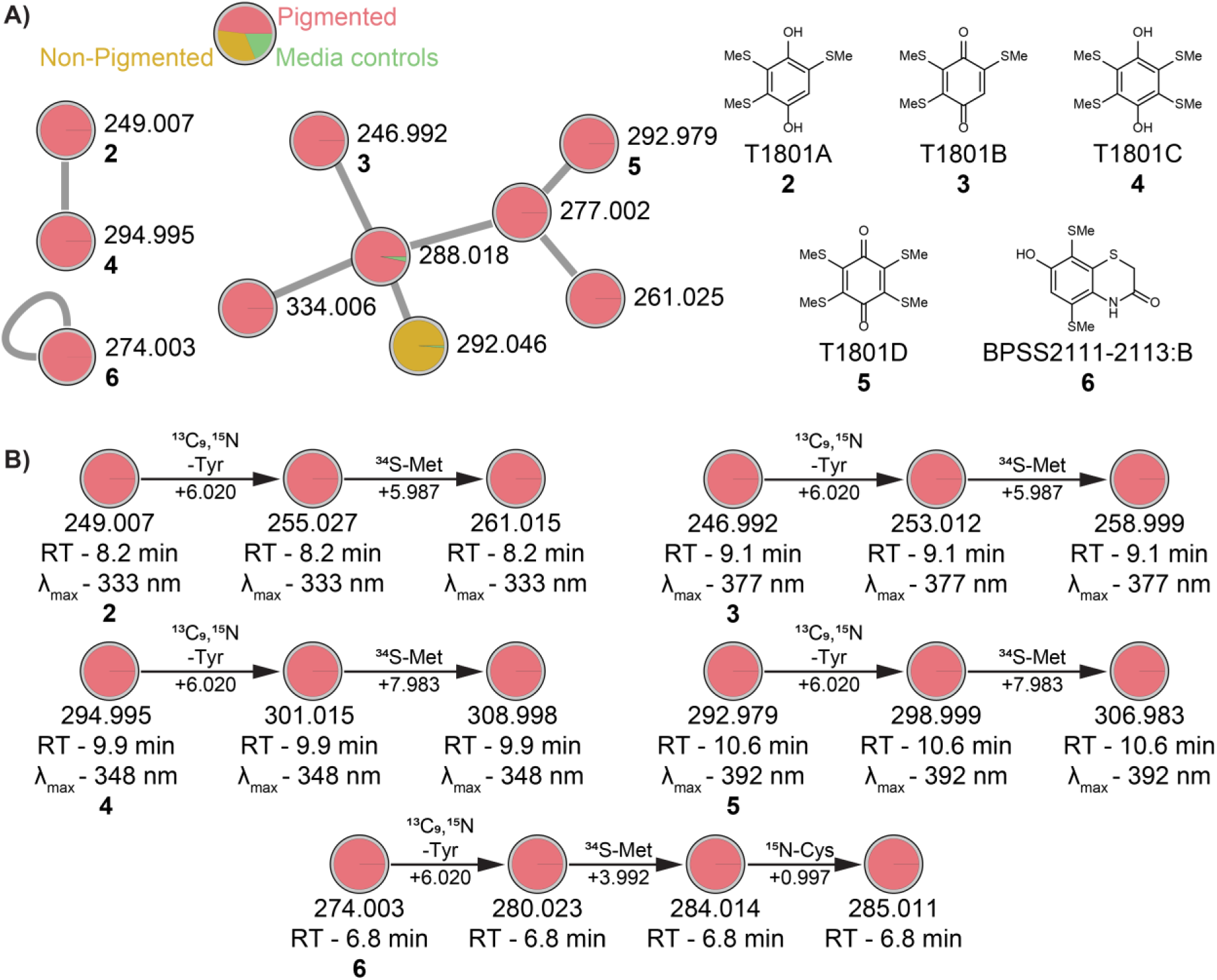
**A**) Molecular network representation of molecular clusters containing nodes for T1801A-D (**2-5**) and BPSS2111-2113:B (**6**). Pie charts mapped onto nodes represent the relative abundance of the feature across each sample type, as indicated by the integrated area under the chromatographic peak (red: pigmented bacterial samples, gold: non-pigmented bacterial samples, green: media controls). **B**) Representation of observed mass shifts after isotope labeling with ^13^C_9_,^15^N-Tyr, ^34^S-Met and ^15^N-Cys. Stable isotope labeling was confirmed by matching MS^2^ spectra (Fig. S12-S17), retention time, and UV-visible absorbance spectra.

The feature with *m/z* 274.003, t_R_ = 6.8 min and predicted molecular formula: C_10_H_11_NO_2_S_3_) was found to have an *m/z* matching that of a metabolite previously reported by Biggins *et. al*.^40^ The authors refer to this compound as BPSS2111-2113:B (**6**) (Fig. 7).^40^ The analysis of metabolomics data acquired on extracts from SILAC experiments supports this annotation where an increase in *m/z* of 6.020 from 274.003 was observed when cultured in ^13^C_9_,^15^N-tyrosine, an additional increase of 3.992 in the presence of ^34^S-methionine, as well as an increase of 0.997 in the presence of ^15^N-cysteine (Fig. 7B, S17). Additional sulfur-containing compounds of related biosynthetic origin were also detected and putatively annotated based on shift of observed *m/z* upon SILAC (Fig. S18, S19).

In summary, untargeted metabolomic analysis of culture extracts from pigmented and non-pigmented *B. cenocepacia* strains as well as genetically modified strains reversed in pigmentation phenotype reveal that pyomelanin production is associated with a distinct profile of secondary metabolite production. Mass spectrometry-based analysis of these metabolites revealed presence of aromatic ring and multiple sulfur atoms, which guided use of appropriate labeled precursors for SILAC. Using SILAC, homogentisic acid is determined as the precursor for biotransformation of the antibiotic trimethoprim and for the biosynthesis of a number of natural products including 3MT-HA (**1**), BPSS2111-2113:B (**2**) and the T1801A-D (**3-6**) family of quinone/hydroquinone antimicrobials in pyomelanin producing *B. cenocepacia*. Production of these compounds involved methylthiolation, the source of which was identified as methionine. Characterization of the biotransformed product of trimethoprim using synthetic methods supported the proposed biochemical pathway for this transformation, which involved methylthiolation of homogentisic acid followed by the addition to trimethoprim. This pathway represents a previously unknown mechanism for bacterial metabolism of the antibiotic trimethoprim, which is specific to the pigmented phenotype and might be widespread among pigmented bacterial pathogens. The discovery of homogentisic acid as a key intermediate for natural product biosynthesis in pigmented strains expand the current knowledgebase of microbial metabolic pathways. Many of the natural products described here have known antimicrobial properties, suggesting that their production can alter the microbial community structure present during the infection. It is now well appreciated that phenotypes capable of altering the microbial community structure present during infection can play an important role in disease progression and development of chronic infection.^41, 42^ These observations advance the current understanding of the chemical makeup of pathogenic *Burkholderia* strains and generate several hypothesis warranting future investigations into the biological functions of the metabolites described in this study. Furthermore, our findings suggest that inhibition of pigment production should be investigated as a strategy to potentiate treatments of bacterial infections by pigment-producing strains with existing antibiotics. Lastly, we propose that the application of computational methods outlined in this study and SILAC is a broadly applicable analytical strategy for discovery of biochemical pathways that are associated with specific microbial phenotypes given the growing availability of isotopically-labelled substrates.

## Methods

### Bacterial strains

All strains used in this study are listed in Table 1. *B. cenocepacia* K56-2 and J2315 strains are clonally related strains belonging to the highly transmissible and epidemic ET-12 lineage.^43–45^ The two wild-type strains were isolated from CF patients, and the laboratory modified strains were provided by Dr. Joanna Goldberg.^9, 44, 46^

### Growth Conditions

Trimethoprim, and sulfadimethoxine were obtained from Sigma-Aldrich. Hexakis(1H,1H,2H-perfluoroethoxy)phosphazene was purchased from SynQuest Laboratories. Salmon sperm DNA and porcine stomach mucin were acquired from Sigma Aldrich and DOPC was purchased from Avanti Polar Lipids. All other media components for M9 minimal medium, LB and SCFM2 synthetic sputum medium were sourced from Fisher Chemical.

SCFM2 media was prepared according to the procedure previously established by Turner and colleagues.^47^ M9 minimal medium was prepared according to the established Cold Spring Harbor protocol, supplementing with 0.4% glucose as a carbon source.^48^ For stable isotope labeling studies, M9 media was additionally supplemented with either 1) L-methionine, and L-tyrosine, 2) L-methionine and L-tyrosine-^13^C_9_,^15^N, or 3) L-methionine-^34^S and L-tyrosine-^13^C_9_,^15^N (400 µM final concentration of methionine and tyrosine). Stable isotope labeling was repeated by supplementing M9 media + 0.4% glucose with either 1) L-cysteine, L-methionine, and L-tyrosine, or 2) L-cysteine-^15^N, L-methionine-^34^S and L-tyrosine-^13^C_9_,^15^N (500 µM final concentration of cysteine, 400 µM final concentration of methionine and tyrosine). For experiments involving ethionine supplementation, M9 media was supplemented with either methionine or ethionine at a final concentration of 620 µM.

For bacterial culturing, frozen glycerol stocks of each strain were aseptically streaked out onto LB-1.5% agar plates before incubating at 37 °C. Once individual colonies were visible, a sterile inoculating loop was used to inoculate 5 mL of LB medium with a single colony from each strain. These cultures were incubated overnight at 37 °C while shaking at 200 rpm and used as starter culture to inoculate three biological replicates of SCFM2 supplemented with 12 mM trimethoprim stock prepared in DMSO to a final concentration of 30 µM, or DMSO to serve as a no-antibiotic control. After inoculating, bacterial cultures and media controls were incubated for 24 h at 37 °C while shaking at 200 rpm.

For all studies in M9 minimal media, culturing was conducted in a similar manner, with the only difference being that cultures were harvested at 48 h rather than 24 h to account for slower growth in minimal media. In addition to the supplemented amino acids, media was further supplemented with either trimethoprim dissolved in DMSO (at a final concentration 30 µM) or an equivalent amount of DMSO as a vehicle control prior to culturing.

### Sample Extraction

Metabolites were extracted from cultures using both liquid-liquid extractions with ethyl acetate (EtOAc) and solid-phase extractions (SPE) using HyperSep™ C18 Cartridges (100 mg bed weight) from Thermo Fisher Scientific. Two separate 5 mL aliquots were taken from each flask, and used for EtOAc and SPE extractions. EtOAc extractions were performed twice with 5 mL of EtOAc each time. After mixing, samples were centrifuged at 2,000×g for 3 min before aspirating off the top organic layer into a scintillation vial with a glass pipette. SPE extractions were performed by first centrifuging cultures at 5,250×g for 10 min and decanting to separate supernatant from cell pellets. Cell-free supernatant was stored at -80 °C before further processing. SPE columns were washed with 5 mL 100% MeCN and equilibrated with 5 mL water before loading with thawed culture supernatant. Analytes were then sequentially eluted from the column using 2.5 mL each of 20%, 50%, and 100% MeCN. SPE elutions were pooled into the scintillation vial containing the corresponding EtOAc extract from the same culture, and pooled EA/SPE extracts were concentrated *in vacuo* before storing at -20 ⁰C.

### Ultra-high-Performance Liquid Chromatography Tandem Mass Spectrometry (UHPLC-MS/MS) Data Acquisition

Dried extracts were dissolved in 150 µL of 80% MeOH (LC-MS grade) containing 1 µM sulfadimethoxine as an internal standard before analyzing 15 µL on a 1290 Infinity II UHPLC system (Agilent Technologies) coupled to an ImpactII ultra-high resolution Qq-TOF mass spectrometer (Bruker Daltonics) equipped with an ESI source. Chromatographic separation was achieved on a Kinetex™ 1.7 µm C18 reversed phase HPLC column (50 × 2.1 mm). Solvent A consisted of water with 0.1% (v/v) formic acid and solvent B consisted of MeCN with 0.1% (v/v) formic acid. The gradient used for chromatographic separation consisted of: 5% B and 95% A for 2 min, linear increase to 95% B over 17 min, hold for 3 min, linear decrease to 5% B in 1 min, and hold for 1 min with a constant flow rate of 0.5 mL/min throughout. For MS spectra acquisition, the instrument was set to capture features from *m/z* 50 - 2000 Da in positive and negative ion mode. External calibration was performed prior to data collection using ESI-L Low Concentration Tuning Mix (Agilent Technologies), with hexakis(1H,1H,2H-perfluoroethoxy)phosphazene employed as internal lock-mass calibrant throughout the run. Ion source parameters were set to 4500 V for capillary voltage, 2 bar for nebulizer gas pressure (N2), 200°C for ion source temperature, 9 L/min for dry gas flow, and a spectral rate of Hz for MS^1^ and 6 Hz for MS^2^. For MS^2^ data acquisition, the eight most intense precursor ions per MS^1^ scan were selected for MS^2^ fragmentation. A basic stepping function was used to fragment ions at 50% and 125% of the CID calculated for each *m/z*, with timing of 50% for each step. Similarly, a basic stepping of collision RF of 550 and 800 Vpp with a timing of 50% for each step and transfer time stepping of 60 and 150 µs with a timing of 50% for each step was employed. The MS/MS active exclusion parameter was set to 2 and the active exclusion was released after 30 sec. The mass of the internal lock-mass calibrant was excluded from MS^2^ acquisition. After each set of triplicates, a wash method was run to clean the column, followed by a solvent blank to monitor for any residual carryover. No carryover was detected in the solvent blanks.

An additional 80 µL of sample extracts were analyzed on a 1260 Infinity UHPLC instrument with a G1315D diode array detector (Agilent Technologies) in series with an AmaZon SL ion trap mass spectrometer (Bruker Daltonics) equipped with an ESI source. Chromatographic separation was performed using a a Kinetex™ 1.7 µm C18 reversed phase HPLC column (50 × 2.1 mm). Solvent A was H_2_O + 0.1% v/v formic acid, and solvent B was MeCN + 0.1% v/v formic acid. The gradient used for chromatographic separation was as follows: 5% B and 95% A from 0– 5 min, linear increase to 100% solvent B from 5–25 min, hold for 5 min, linear decrease to 5% B from 30-31 min, hold for 2 min, linear increase to 100% B from 33-34 min, hold for 4 min, linear decrease to 5% B from 38-39 min, and a final hold for 3 min with a constant flow rate of 0.5 mL/min throughout the run. Mass spectrometry data was acquired in positive ionization mode in the mass range m/z 100–2200 Da. Ion source parameters were set to 4500 V for capillary voltage, 5 psi for nebulizer gas pressure (N2), 150°C for ion source temperature, and 3 L/min for dry gas flow. For MS^2^ data acquisition, the two most intense precursor ions per MS^1^ scan were selected for MS^2^ fragmentation. Annotations for T1801A-D (**2-5**) were supported by comparing their experimental UV spectra with values reported in prior literature.^39^

### Data Processing, Feature-based Molecular Networking, and Feature Annotation

Raw LC-MS/MS spectral data in Bruker format (.d) were converted to centroided, lock-mass corrected format (.mzXML) in Bruker® Compass DataAnalysis using a “Process with Method” batch script. The resulting .mzXML files were subsequently uploaded to the UCSD MassIVE online repository and made publicly available under the ID MSV000089255. Converted spectral files were uploaded to MZmine2 (v.2.5.1) for feature detection.^17^ Data files were batch processed and filtered using a MS^1^ mass detection signal threshold of 5.0 × 10^2^ counts. The following parameters were applied: (i) chromatogram builder (minimum time span: 0.05 min; minimum intensity of the highest data point in the chromatogram: 1.5 × 10^3^; *m/z* tolerance: 10 ppm); (ii) chromatogram deconvolution (local minimum search, *m/z* range for MS^2^ scan pairing: 0.025 Da; retention time range for MS^2^ scan pairing: 0.2 min); (iii) isotopic peaks grouper (*m/z* tolerance: 10 ppm; retention time tolerance absolute: 0.1 min; maximum charge: +3; monotonic shape: true; representative isotope: most intense); (iv) join aligner (*m/z* tolerance: 10 ppm; retention time tolerance: 0.1 min); (v) feature list rows lifter (minimum peaks in a row: true, 2; minimum peaks in an isotope pattern: true, 2; keep only peaks with MS^2^ scan (GNPS): true; reset the peak number ID: true); (vi) duplicate peak filter (filter mode: new average; *m/z* tolerance: 5 ppm; t_R_ tolerance: 0.1 min); (vii) peak finder (intensity tolerance: 10%; retention time tolerance (absolute): 0.1; m/z tolerance: 10 ppm). A table with ion intensities for each feature across each sample was exported (.csv format) for statistical analyses and the “Export for SIRIUS” module was used to generate an .mgf file for batch analysis with SIRUS (v.4.8.2). SIRIUS 4 with CSI:FingerID was used to predict molecular formulas for these features.^49, 50^ CANOPUS was also employed to predict compound classes based on ClassyFire ontology for each feature, providing biological insight even when structures could not be predicted.^51, 52^ Additionally, the “Export for GNPS” module was used to convert and export the feature quantification table (.csv format) and the corresponding list of MS^2^ spectra linked to MS^1^ features (.mgf format) needed to generate a feature-based molecular network. The feature quantitation table and .mgf file were submitted to the Global Natural Products Molecular Networking (GNPS) platform along with a metadata file (.txt format) containing sample information, and a feature-based molecular network was created.^19^ Molecular networks and parameters used are accessible at the following links: https://gnps.ucsd.edu/ProteoSAFe/status.jsp?task=d5b3c60078da4ea08505439a072780d0 (positive mode network) and https://gnps.ucsd.edu/ProteoSAFe/status.jsp?task=bb779c06162d4cef98c819a608d08823 (negative mode network). A parent mass tolerance of 0.01 Da and a MS/MS fragment ion tolerance of 0.01 Da were applied to create consensus spectra. The network was created with the edges filtered to have a cosine score above 0.7 and at least 4 matched peaks, and edges connecting two nodes were set to be kept in the network if each of the nodes appeared in each other’s respective top 10 most similar nodes. All the mass spectra in the generated network were queried against the spectral libraries available on GNPS. Library spectra were considered to be matches to experimental spectra if spectra contained more than 4 matched peaks with a cosine similarity score above 0.7. Networks were exported to Cytoscape for visualization, and nodes appearing in media or solvent blanks were removed for clarity (unless otherwise indicated).^53^ Next, we implemented MS2LDA analysis separately on each network to recognize recurring patterns of common MS^2^ fragments and neutral losses detected in our dataset, which correspond to molecular substructures. MS2LDA analysis and the parameters used are accessible at the following links: https://gnps.ucsd.edu/ProteoSAFe/status.jsp?task=827d4d51ca5741b2b4db3b4a55ac3bc6 (positive mode network) and https://gnps.ucsd.edu/ProteoSAFe/status.jsp?task=3a9d184119894782a187cb0b5384c3d8 (negative mode network).

To filter out duplicate features corresponding to compounds detected in both positive and negative ion mode, the MScombine R package was used combine data into a single feature quantification table.^18^ The UpSet plot was generated with the UpSetR package, using the merged polarity feature table as input.^21^ In the UpSet plot, samples were grouped together on the basis of their pigment phenotype and exposure to trimethoprim. To identify metabolomic features which distinguish between pigmented and non-pigmented *B. cenocepacia* strains, the merged polarity feature table was uploaded to statistical analysis on the MetaboAnalyst web server.^54^ Samples were grouped on the basis of strain and trimethoprim exposure. Group 1 consisted of *B. cenocepacia* J2315 WT samples without trimethoprim, group 2 was *B. cenocepacia* J2315 WT samples with trimethoprim, group 3 was *B. cenocepacia* K56-2 J*hmgA* without trimethoprim, group 4 was *B. cenocepacia* K56-2 J*hmgA* samples with trimethoprim, group 5 was *B. cenocepacia* J2315 K*hmgA* samples without trimethoprim, group 6 was *B. cenocepacia* J2315 K*hmgA* samples with trimethoprim, group 7 was group 5 was *B. cenocepacia* K56-2 WT samples without trimethoprim, and group 8 was *B. cenocepacia* K56-2 WT samples with trimethoprim. Data was subjected to log transformation prior to conducting PCA and PLS-DA analysis. For the PLS-DA model, the “class order matters” option was selected. The leave one out cross-validation (LOOCV) method was used for cross-validation of the three-component PLS-DA model, yielding an R^2^ value (goodness of fit) of 0.998 and Q^2^ value (goodness of prediction) of 0.820. VIP scores were calculated as a measure of a feature’s importance in the PLS-DA model, and features with higher VIP scores were prioritized for structural annotation. To visualize metabolites differentially detected between pigmented and non-pigmented samples onto loadings plots, the same dataset was re-uploaded to MetaboAnalyst and samples grouped on the basis of pigment presence. Fold change analysis and two-tailed t-tests were performed to identify differentially detected metabolites, using an FDR adjusted p-value threshold of 0.05.

Literature searching for compounds known to be produced by *Burkholderia* spp. and other closely related bacteria was conducted to aid with annotating features of interest not identified via GNPS spectral library searching. Molecules that were flagged as potential candidates for unknown features were confirmed by manually inspecting raw MS^2^ spectra and either comparing to published MS^2^ spectra when available, or against MS^2^ spectra acquired from commercially purchased analytical standards. The BLASTP algorithm was used to search *B. cenocepacia* J2315 (NCBI taxid: 216591) and K56-2Valvano (NCBI taxid: 985076) taxonomy databases for candidate protein sequences exhibiting statistically significant similarity to L-methionine γ-lyase from *P. putida* (UniProtKB accession: P13254) and BPSS2111 from *B. pseudomalleli* K96243 (UniProtKB accession: Q63IG2) with default algorithm parameters.^55^ Positive and negative ionization mode datasets were analyzed in two separate batches using SIRIUS (v.4.8.2) integrated with CSI:FingerID and CANOPUS.^49, 50, 52, 56^ SIRIUS 4 was implemented to predict molecular formulas and develop fragmentation trees to aid with manual annotation of MS^2^ spectra, using the default settings for a qTOF instrument.^49, 56^ CSI:FingerID was applied to predict structural fingerprints of unknown features, which were then queried against all available chemical databases to provide a ranked list of possible known structures.^50^ CANOPUS was deployed to classify features into ClassyFire compound classes, providing biological insight into detected compounds eluding structural annotation.^51, 52^

### Headspace Detection and Analysis

Volatile headspace analysis was performed using vacuum assisted sorbent extraction (VASE) coupled with a gas chromatography mass spectrometer (GC-MS), according to previously established methods.^57^ Briefly, samples (methanethiol standard (30 µL of 2 mM), growth medium (200 µL, 24 h), bacterial cultures (200 µL, 24 h of growth)) and pre-cleaned type 1 glass vials (VOA; Thermo Scientific) were chilled on ice for 10 min. A VASE Sorbent Pen cartridge containing Tenax was placed into each vial containing sample and held in place via a lid liner. Air was evacuated from vials using a vacuum pump before placing vials in a shaking incubator for 1 h at 70 ⁰C. After extracting, vials were placed on a metal block equilibrated at −20 °C for 15 min to remove water from the headspace. VASE pens were removed from the vials and their contents were run on an Agilent GC-MS (7890A GC and 5975C inert XL MSD with Triple-Axis Detector) with a DB-624 column. The Entech GC/MS method has a 38 min runtime with a pre-heat duration for 2 min at 260 °C, desorption duration for 2 min at 260 °C, bake-out duration for 34 min at 260°C, and post-bake duration for 2 min at 70 °C. Analysis and quantification of peaks were performed in the Agilent ChemStation software.

### Synthesis Reactions

All synthesis reactions were performed using chemicals and solvents from Thermo Fisher Scientific™, Spectrum™ Chemical, and Cayman Chemical. 3MT-HA (**1**) was synthesized by adding homogentisic acid (1.0 equiv) and sodium methanethiolate (10.0 equiv) to a sealed, oven-dried round-bottom flask (Fig. 6) under ambient atmospheric conditions. MeOH (5 mL solvent per mmol homogentisic acid) was slowly added. The reaction mixture was heated to 65⁰C in an oil bath for 1 h, stirring continuously. The resulting reaction mixture displayed an amber yellow color. Two volumes H_2_O were added to the mixture, before extracting thrice with two volumes EtOAc. The organic layers were pooled and concentrated *in vacuo* to yield a light brown solid. **HRMS** (ESI-qToF) *m/z* calculated for C11H15O4S3 ([M+H]^+^): 307.013, observed 307.012. **UV λ_max_** in 80% MeOH: 334 nm.

The trimethoprim biotransformation product with *m/z* 551.145 was prepared by dissolving trimethoprim (14.5 mg, 1 equiv) in 2.5 mL DMSO and activated to a reactive iminoquinone methide intermediate by addition of 75 µL sodium hypochlorite (5% available chlorine) followed by heating at 65⁰C for 20 min while stirring continuously.^34^ Next, triethylamine (7 µL, 1 equiv) was added to reaction mixture. Finally, 3.2 mg of the crude solid produced during synthesis of **1** dissolved in 2.5 mL DMSO was added, followed by heating at 65⁰C for 1 h while stirring continuously. Two reaction volumes of H_2_O were added to the mixture, before extracting thrice with two reaction volumes EtOAc. The organic layers were pooled and concentrated *in vacuo* via rotary evaporation to yield a light brown solid. **HRMS** (ESI-qToF) *m/z* calculated for C24H31O5S3 ([M+H]^+^): 551.145, observed 551.144. **UV λ_max_** in 80% MeOH: 272 nm.

## Supporting Information

PCA Loadings; PLS-DA scores plot; mass spectral analysis of transformation products of trimethoprim; adenine and anthranilic acid; extracted ion chromatograms of recurring substructure; trimethylthiolated homogentisic acid; MS^2^ spectra of labeled and unlabeled trimethylthiolated homogentisic acid; GC/MS analysis for methanethiol; MS^2^ spectra of labeled and unlabeled biotransformed trimethoprim; spectral comparison of synthesized and bacterially produced trimethylthiolated homogentisic acid; biotransformed trimethoprim; MS^2^ spectra of labeled and unlabeled natural products T1801 A; T1801 B; T1801 C; T1801 D; BPSS2111-2113:B; *m/z* 286.020; *m/z* 301.997.

## AUTHOR INFORMATION

### Corresponding Author

Neha Garg − School of Chemistry and Biochemistry, and Center for Microbial Dynamics and Infection, Georgia Institute of Technology, Atlanta, Georgia 30332-2000, United States; orcid.org/0000-0002-2760-7123;

### Author Contributions

The manuscript was written through contributions of all authors. All authors have given approval to the final version of the manuscript.

## Acknowledgements

We thank Y. Yang (Georgia Institute of Technology) for his contribution in designing synthesis of methylthiolated homogentisic acid. We also thank K. Whiteson (University of California Irvine) for sharing her expertise in application of the VASE method for overhead space analysis by GC-MS and providing access to the instrumentation in her laboratory for acquisition of GC-MS data.

## Abbreviations

UHPLC-HRMS/MS: ultra-high performance liquid chromatography high-resolution tandem mass spectrometry
3MT-HA: trimethylthiolated homogentisic acid.

## Supporting Information

**Figure S1.**
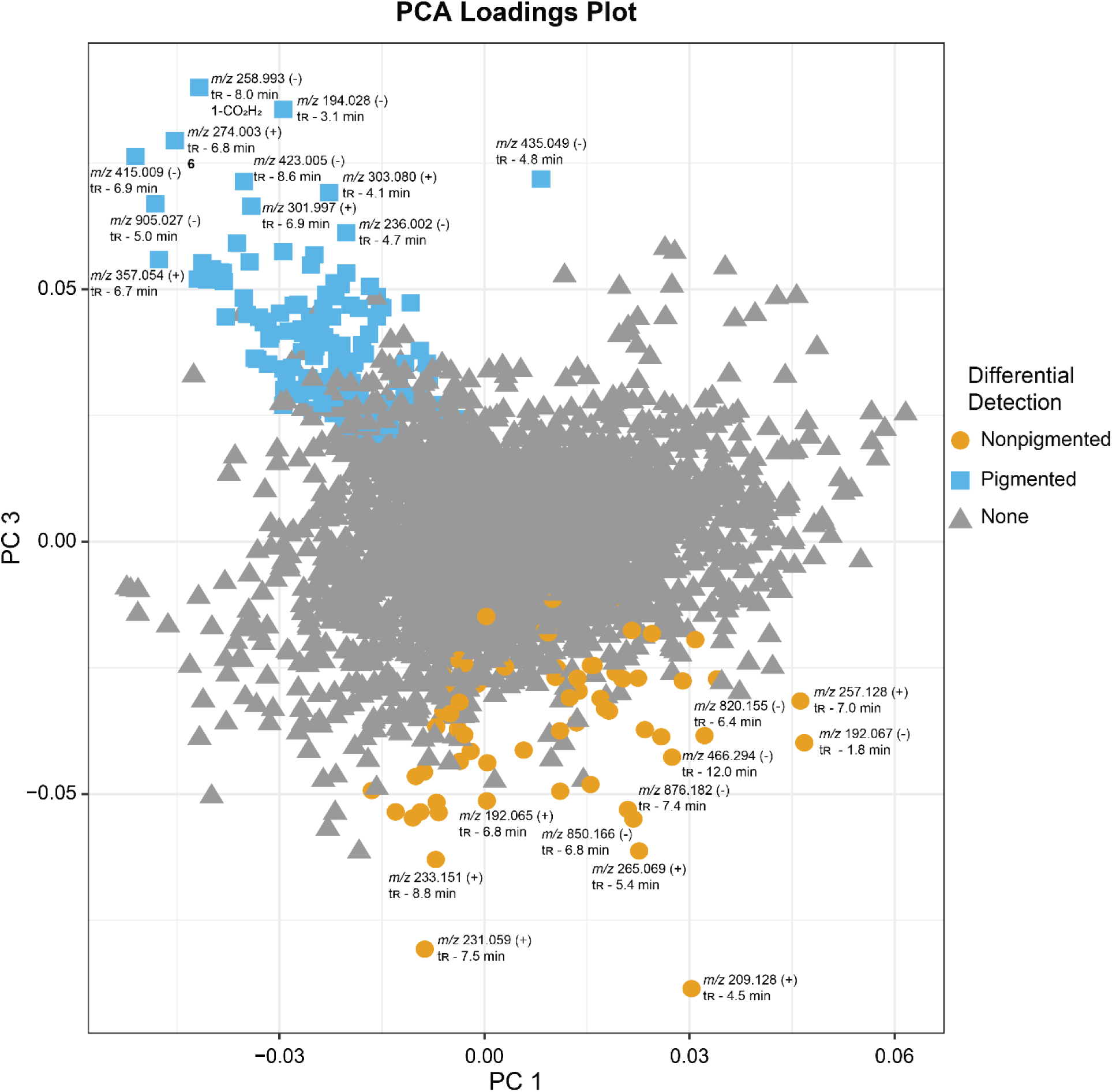
PCA Loadings Plots for PC1 and PC3 generated using on MetaboAnalyst. A t-test (performed on MetaboAnalyst) was used to identify metabolites differentially detected (p_adj_ < 0.05) in pigmented/non-pigmented samples. Features detected at significantly higher abundances in pigmented samples are indicated by blue squares, while features detected at higher abundances in non-pigmented samples are indicated by orange circles.

**Figure S2.**
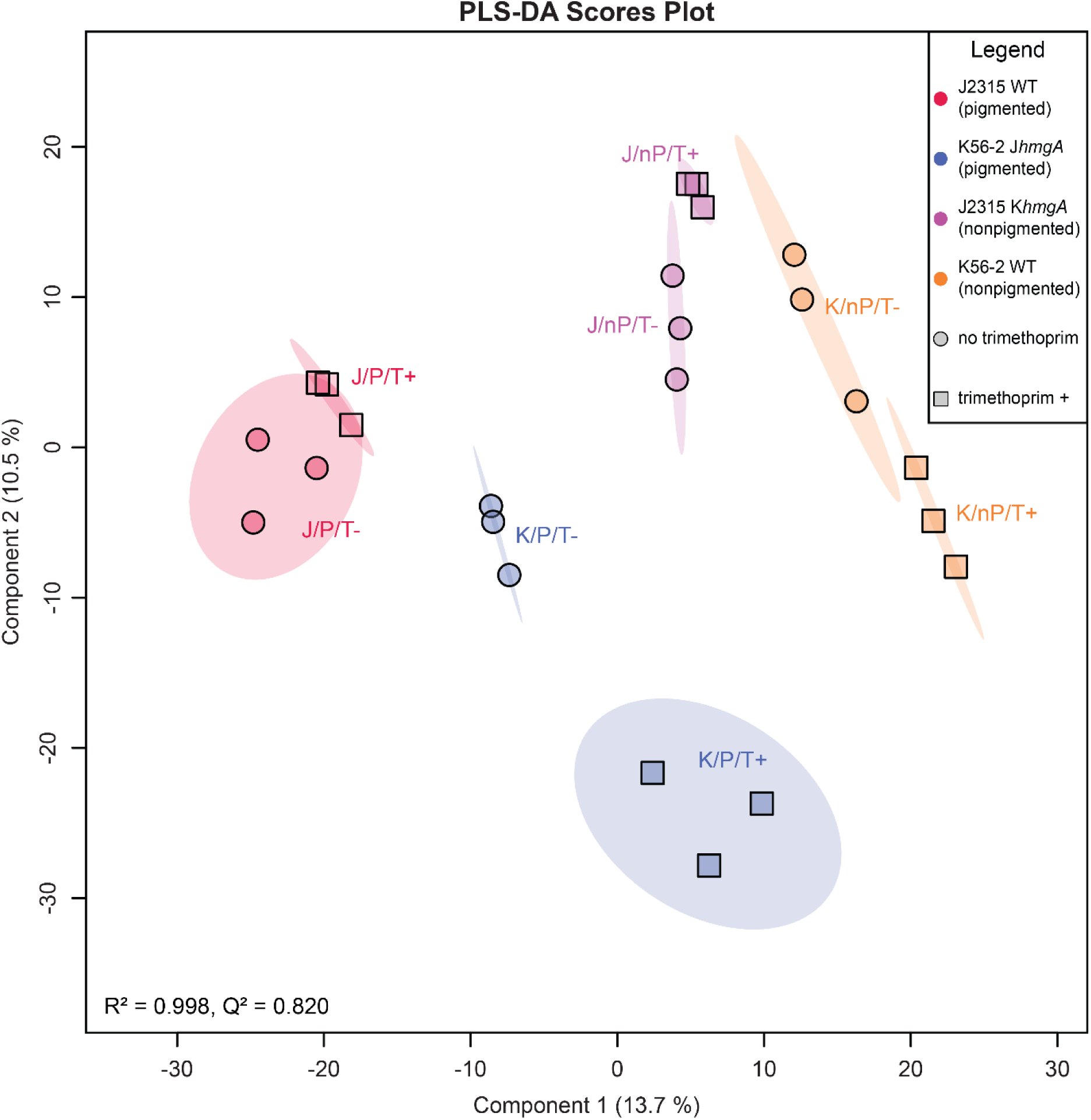
PLS-DA scores plot generated using MetaboAnalyst. Note that while a three component PLS-DA model was used for feature selection and cross-validation, only the first two components are plotted for clarity. Cross-validation was performed using a leave one out cross-validation (LOOCV) approach, yielding an R^2^ value (goodness of fit) of 0.998 and Q^2^ value (goodness of prediction) of 0.820 appropriate for this method.

**Figure S3.**
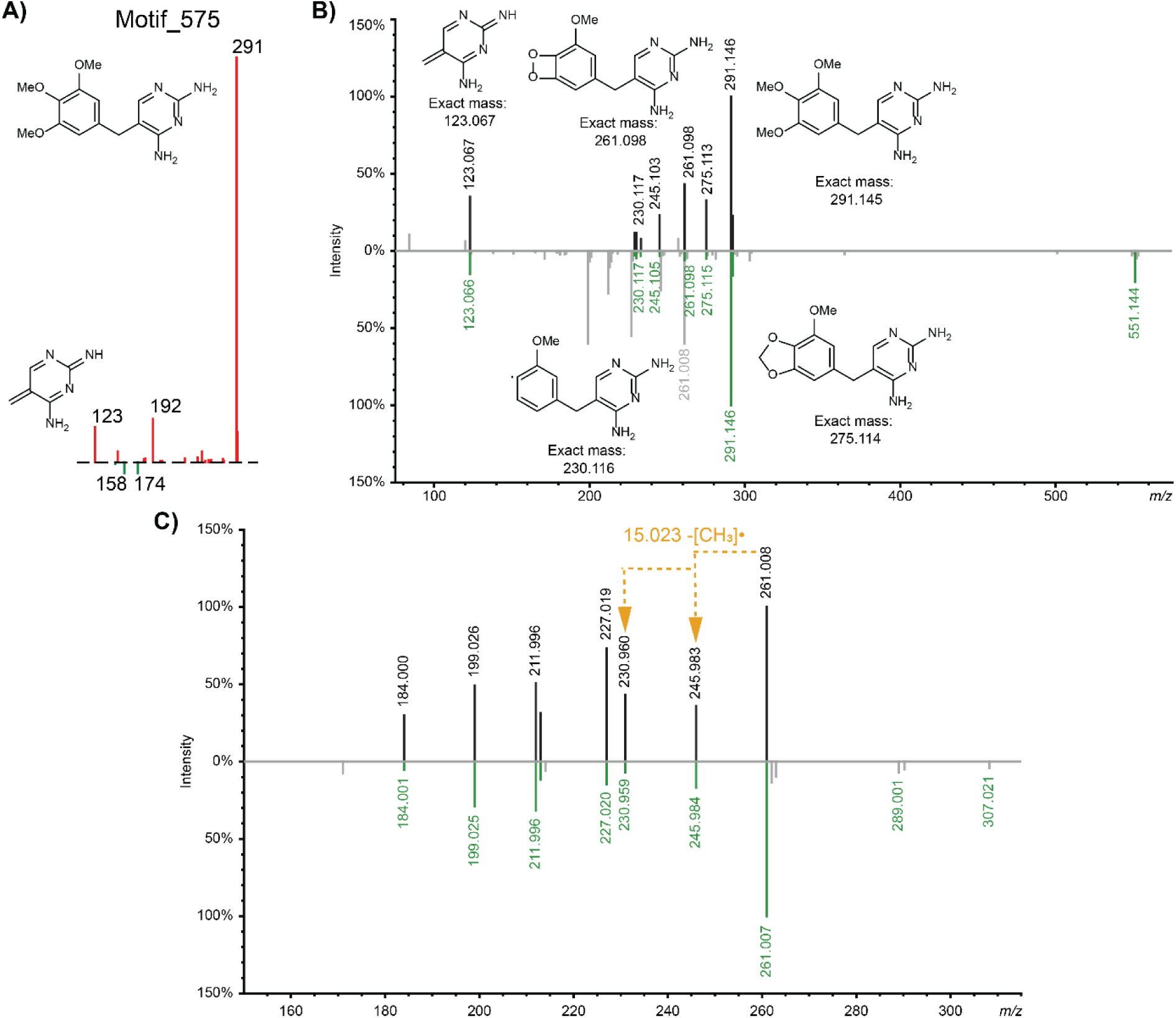
A) Mass2Motif 575 corresponding to trimethoprim substructure. B) Mirror plots comparing the MS^2^ spectra of trimethoprim (top) and metabolite feature with *m/z* 551.145.C) Mirror plots comparing the MS^2^ spectra of metabolite features with *m/z* 261.007 (top) and *m/z* 307.021 (bottom).

**Figure S4.**
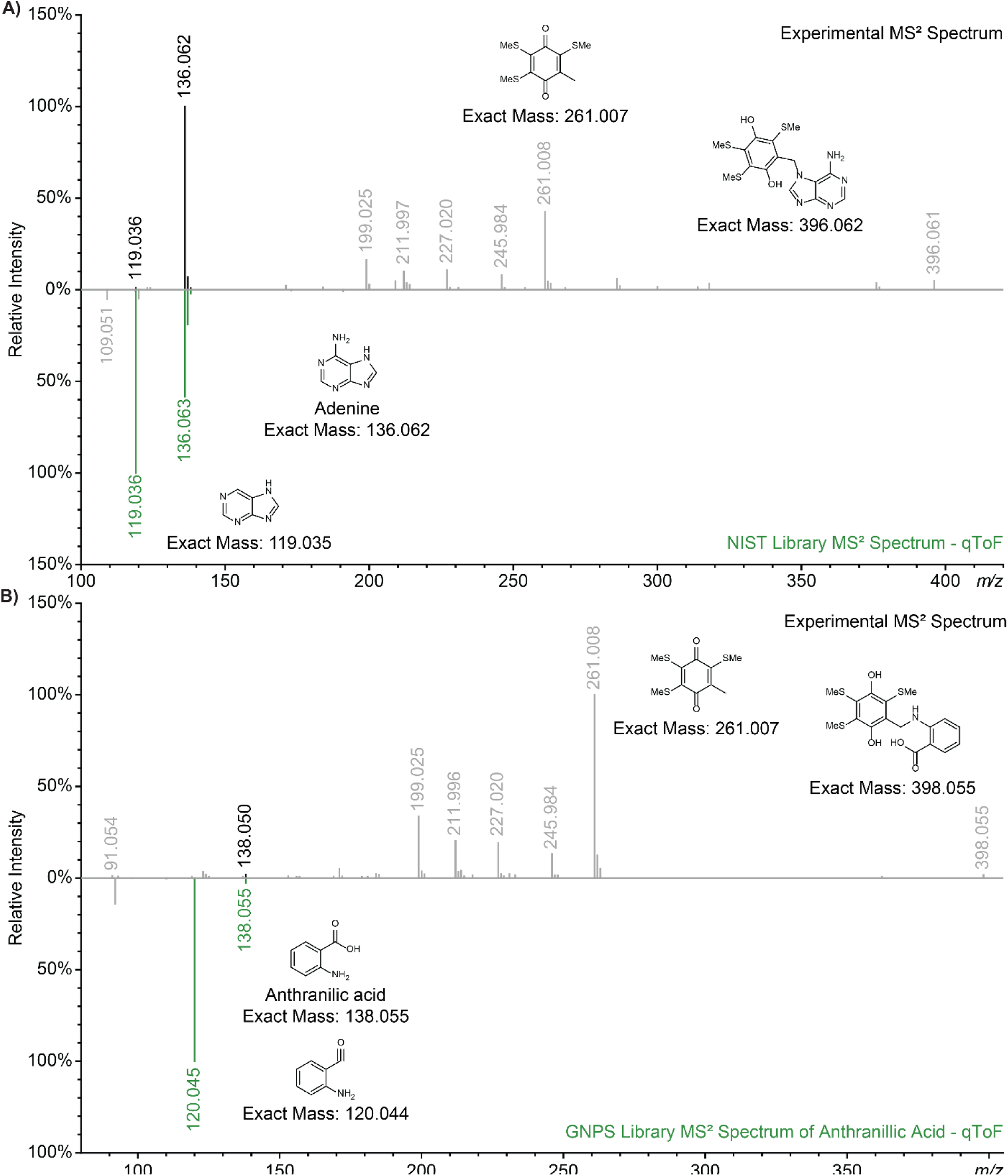
A) Mirror plot comparing the experimental MS^2^ spectrum obtained from *m/z* 396.061 (261.007+ with the spectrum available for adenine in the NIST Tandem Mass Spectral Library. B) Mirror plot comparing the experimental MS^2^ spectrum obtained from *m/z* 398.055 with a spectrum available for anthranilic acid in the GNPS spectral database. These experimental MS^2^ spectra also share fragments with 1 (*m/z* 261.008, *m/z* 245.984, *m/z* 227.020, *m/z* 211.997, *m/z* 199.025), leading to their annotation as biotransformation products.

**Figure S5.**
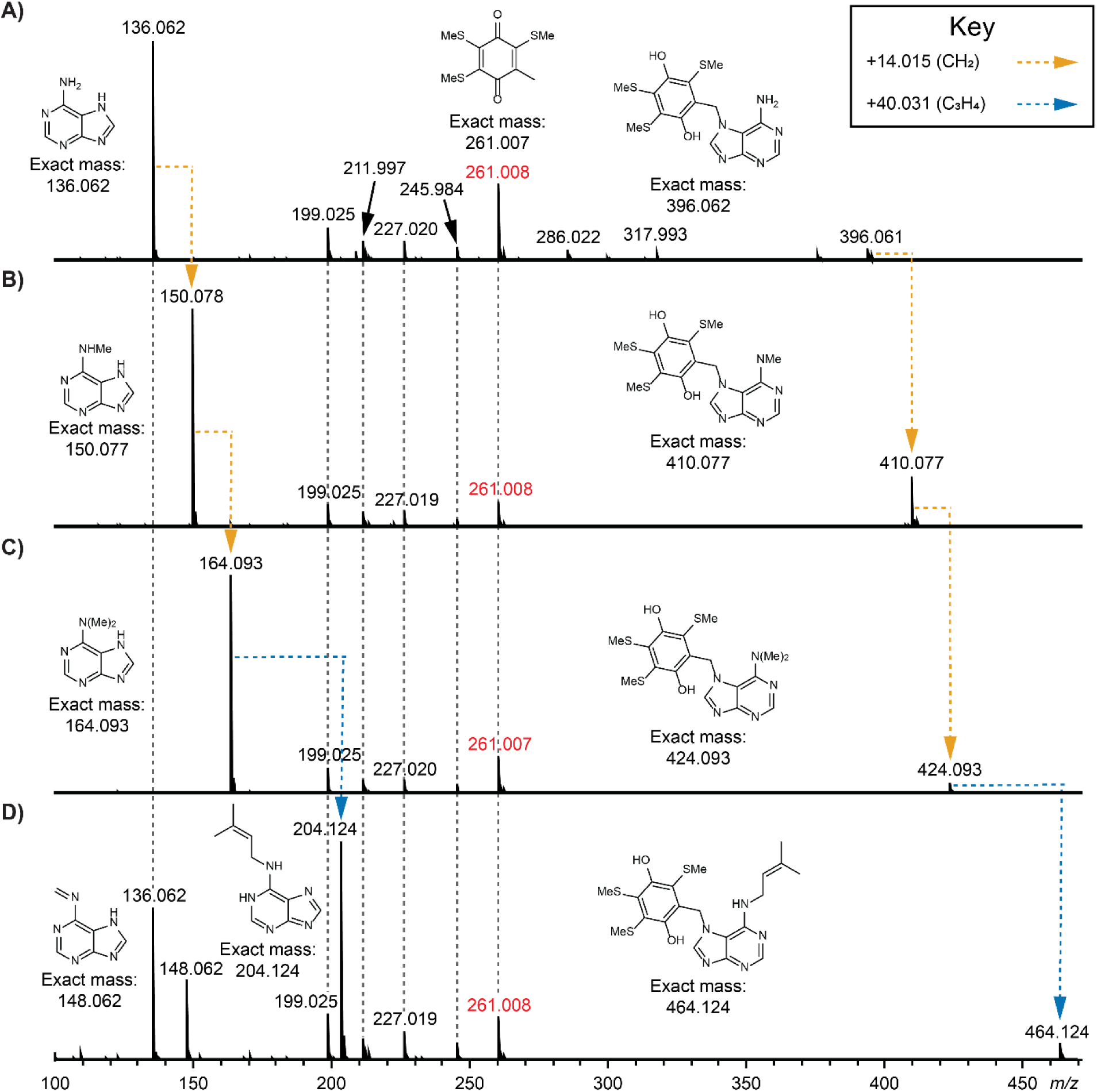
Annotated MS^2^ spectra supporting annotations for biotransformation products of A) adenine, B) methyladenine, C) dimethyladenine, and D) isopentenyladenine. Note that while mass spectrometry cannot conclusively determine the exact location of the additional mass gain, the *N*6 position was deemed most likely based on established biological precedence.

**Figure S6.**
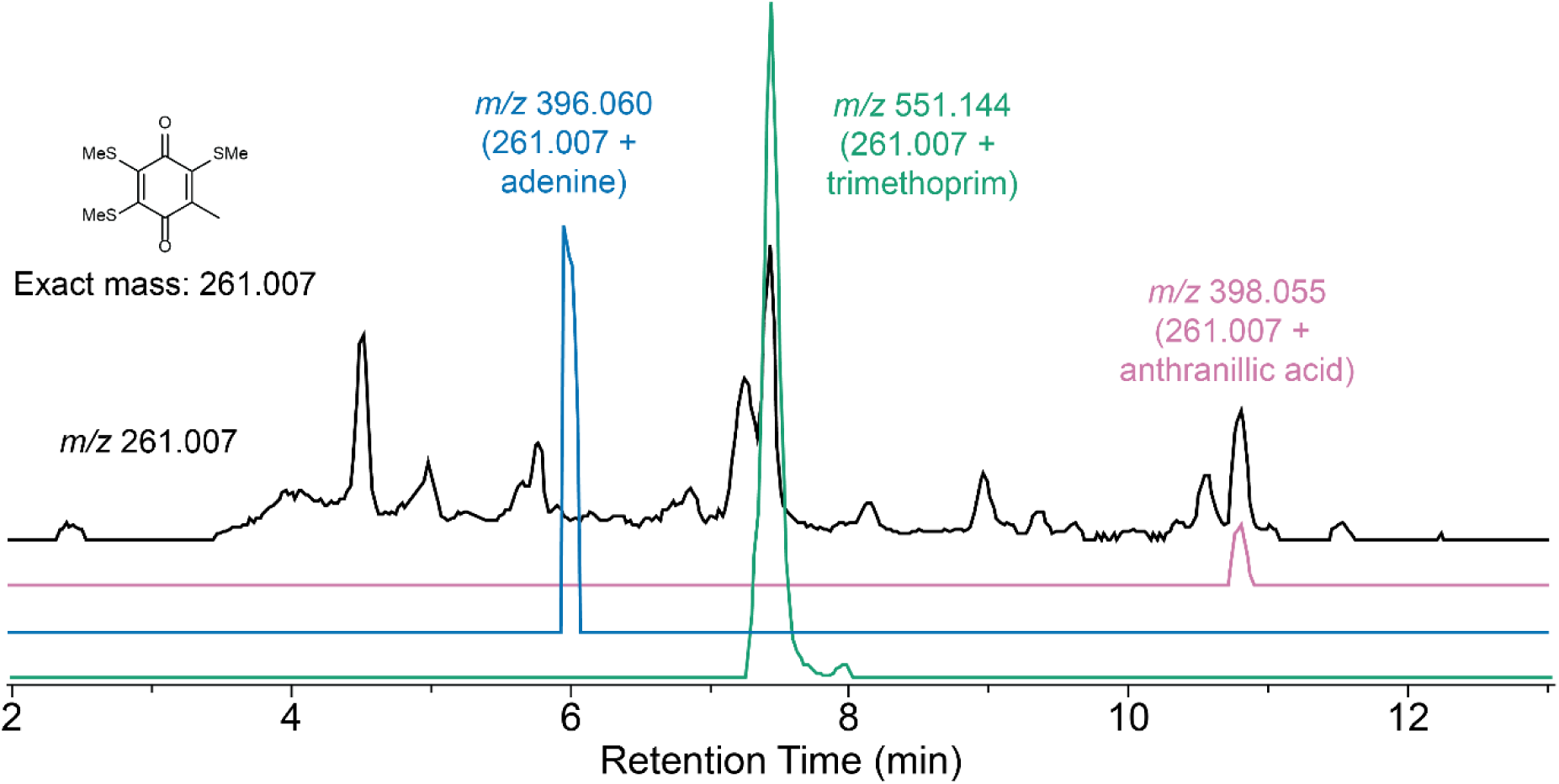
Extracted ion chromatogram (EICs) of the recurring substructure with *m/z* 261.007 (black), as well as adenine (blue), trimethoprim (green) and anthranilic acid (purple) biotransformation products in positive ionization mode (data is shown for the wild-type *B. cenocepacia* J2315). At each peak, the substructure with *m/z* 261.007 is detected as an in-source fragment of a metabolite with higher mass detected at the same retention time. EIC peak heights are not drawn to scale for the sake of clarity.

**Figure S7.**
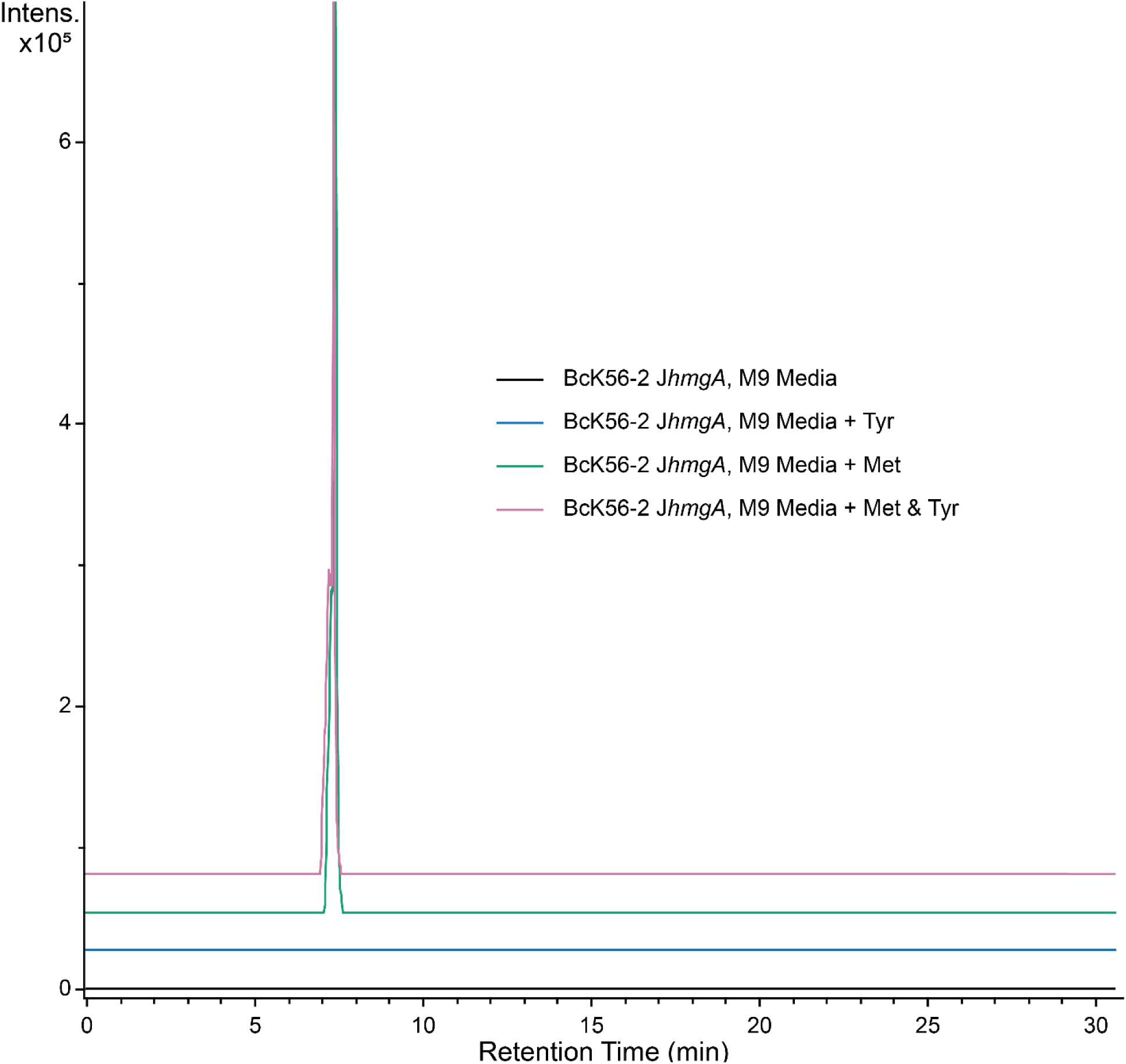
Extracted ion chromatogram (EICs) of *m/z* 307.013 (1) detected in extracts of *B. cenocepacia* K56-2 J*hmgA* cultured in M9 minimal media (black trace), M9 + tyrosine (blue trace), M9 + methionine (green trace), and M9 + methionine & tyrosine (purple trace).

**Figure S8.**
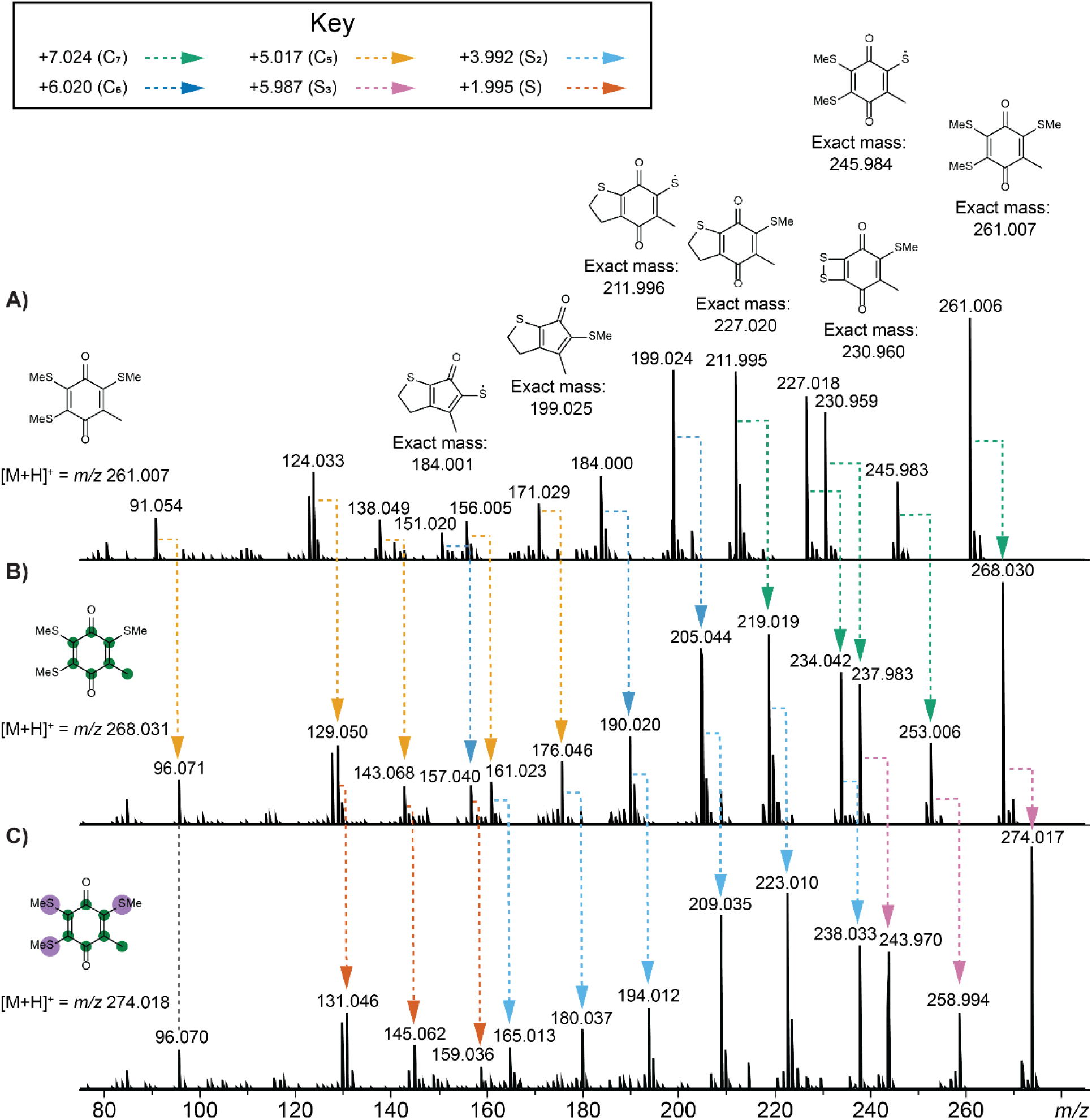
Annotated MS^2^ spectra of the recurring in-source fragment corresponding to the reactive substructure derived from trimethylthiolated homogentisic acid when grown in M9 minimal media supplemented with A) L-methionine and L-tyrosine, B) L-methionine and L- tyrosine-^13^C_9_,^15^N, or C) L-methionine-^34^S and L-tyrosine-^13^C_9_,^15^N.

**Figure S9.**
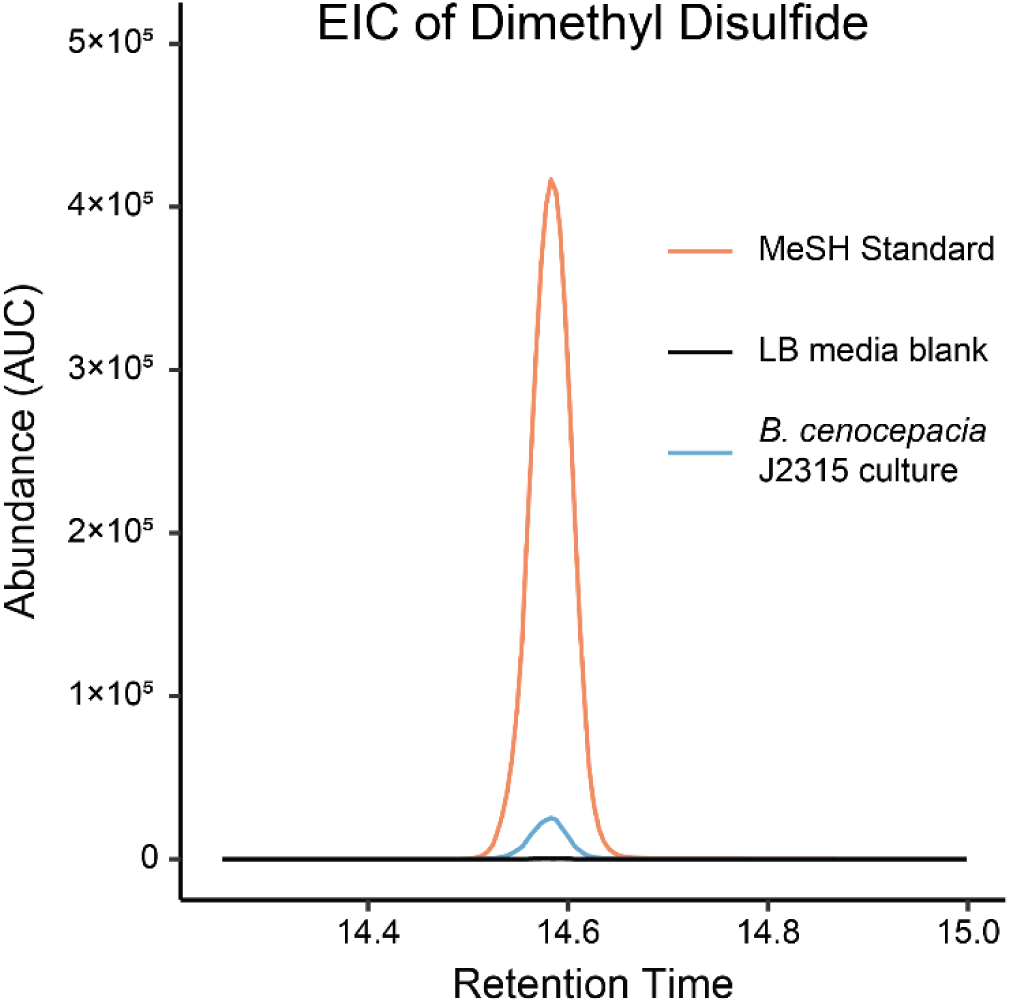
Extracted ion chromatogram of dimethyl disulfide (formed by oxidative dimerization of methanethiol) obtained by performing headspace GC/MS analysis on a commercial methanethiol standard, *B. cenocepacia* J2315 culture, and LB media blank sample.

**Figure S10.**
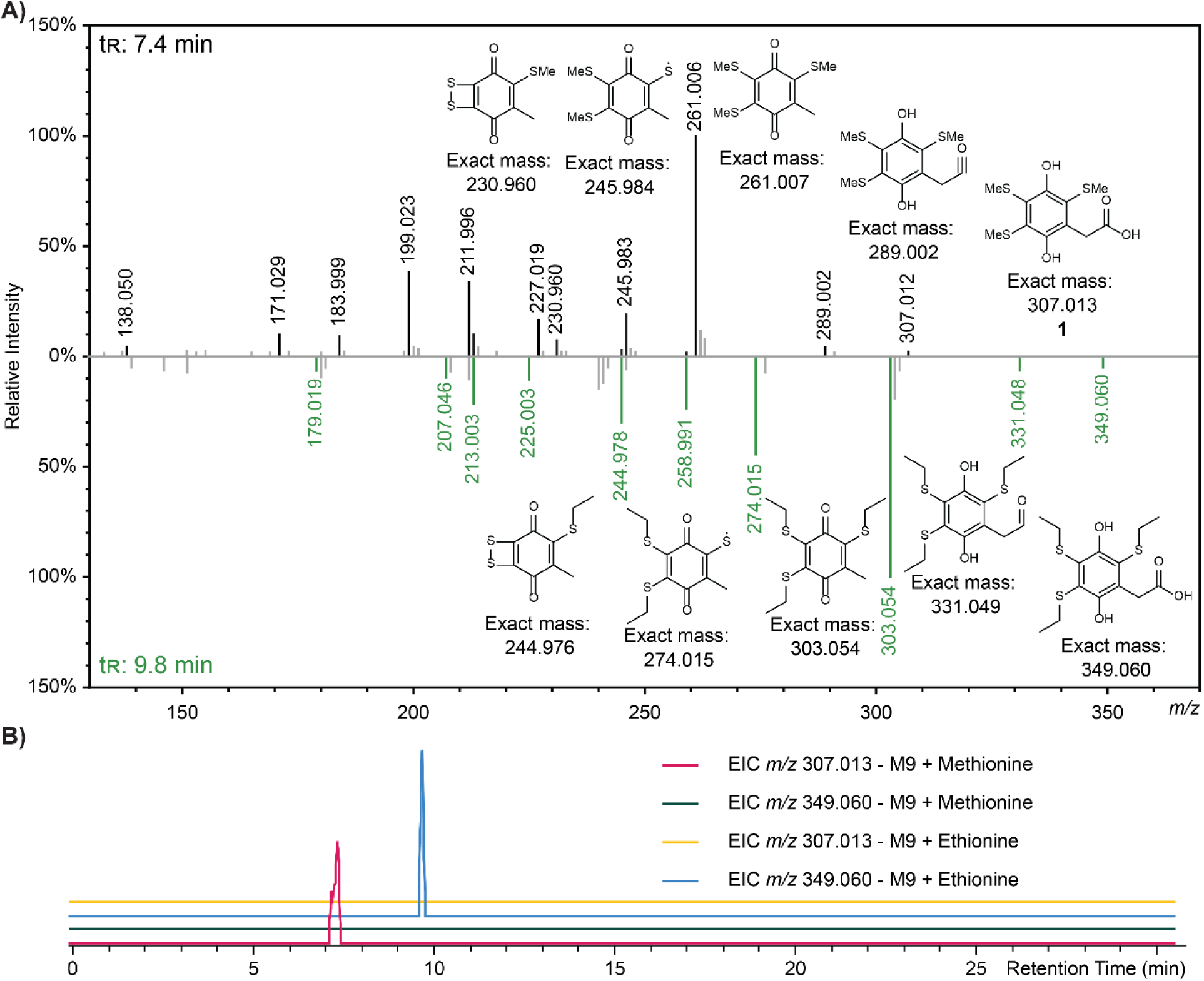
A) Mirror plot of MS^2^ spectrum of trimethylthiolated homogentisic acid (1, top) detected in extracts of *B. cenocepacia* J2315 wild-type cultured in M9 minimal media supplemented with methionine and MS^2^ spectrum of ethylthiolated analog (2-(2,4,5- tris(ethylthio)-3,6-dihydroxyphenyl)acetic acid, bottom) detected in extracts of *B. cenocepacia* J2315 wild-type cultured in M9 minimal media supplemented with ethionine. **B)** Extracted ion chromatograms of **1** and ethylthiolated analog with *m/z* 349.060 in extracts of *B. cenocepacia* J2315 wild-type cultured in M9 minimal media supplemented with either methionine or ethionine.

**Figure S11.**
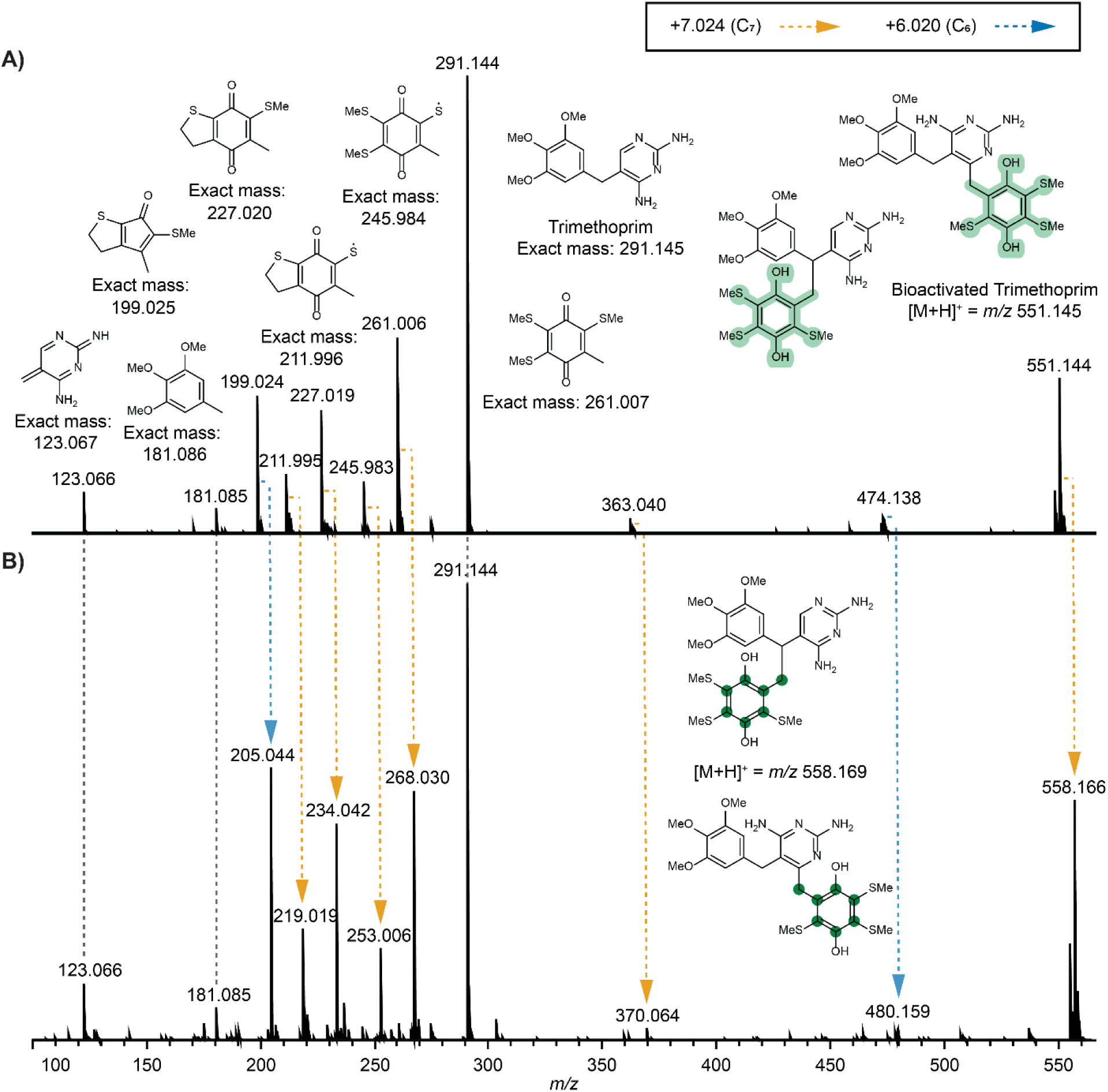
Annotated MS^2^ spectra of the bioactivated trimethoprim product. The data is shown for extracts of bacterial cultures grown in M9 minimal media supplemented with trimethoprim as well as either A) L-methionine and L-tyrosine or B) L-methionine and L-tyrosine-^13^C_9_,^15^N.

**Figure S12.**
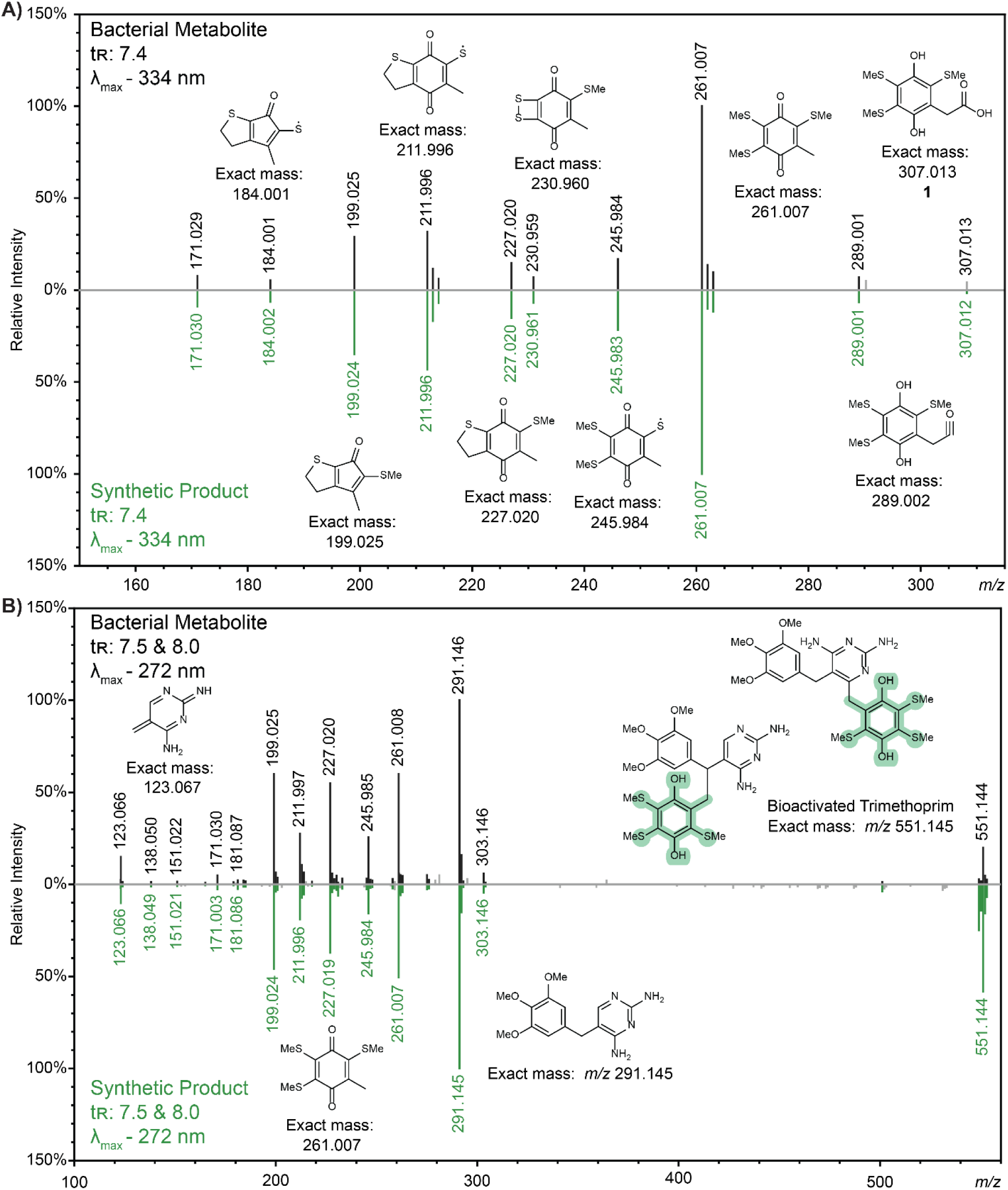
The MS^2^ spectral, retention time, and UV-visible absorbance spectral match of bacterially produced and the synthetic compounds A) 1, and B) the trimethoprim biotransformation products with *m/z* 551.144.

**Figure S13.**
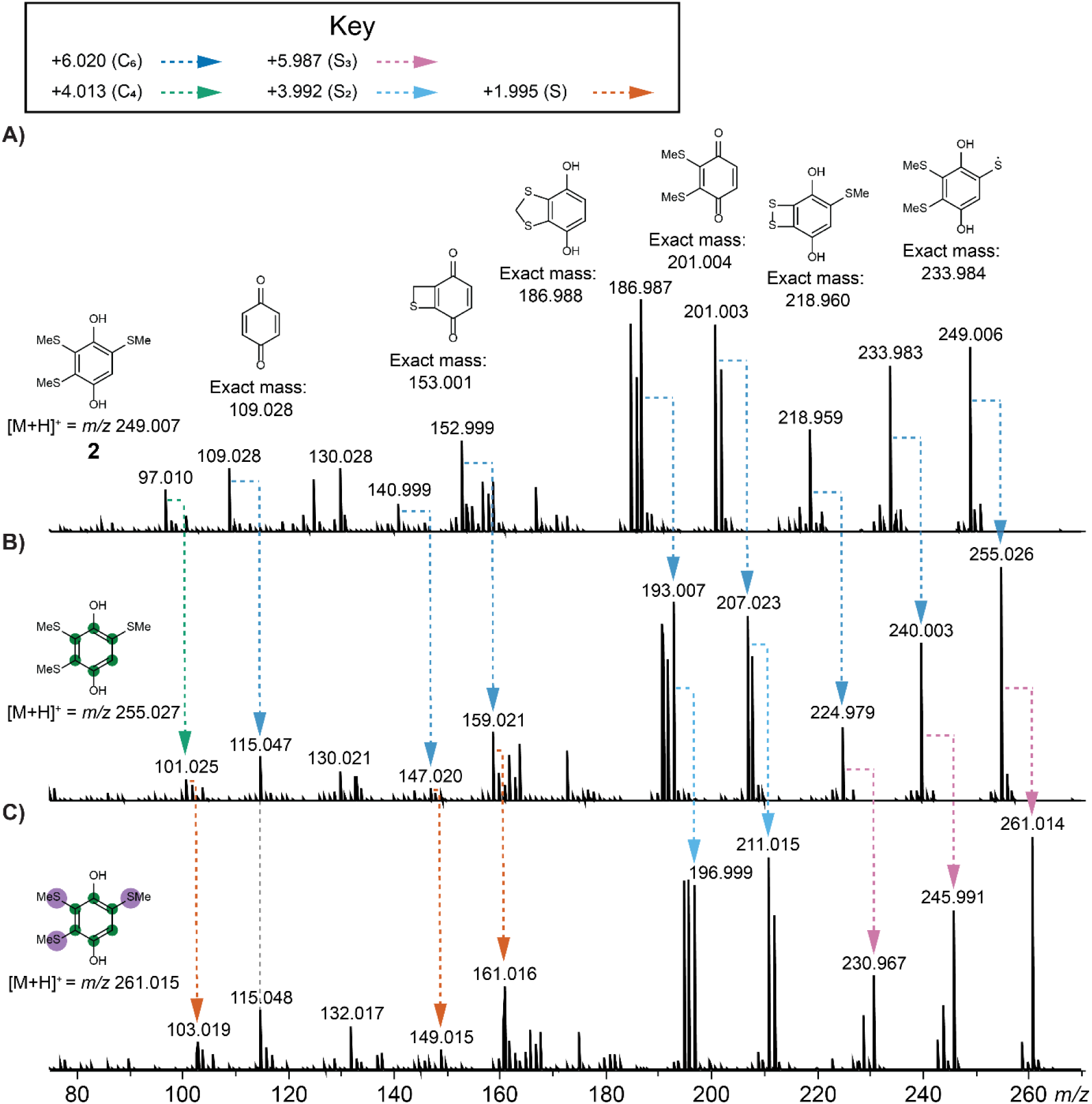
Annotated MS^2^ spectra highlighting mass shifts due to isotope labeling of T1801 A (2) when bacterial strains were cultured in M9 minimal media supplemented with A) L-methionine and L-tyrosine, B) L-methionine and L-tyrosine-^13^C_9_,^15^N, or C) L-methionine-^34^S and L-tyrosine- ^13^C_9_,^15^N.

**Figure S14.**
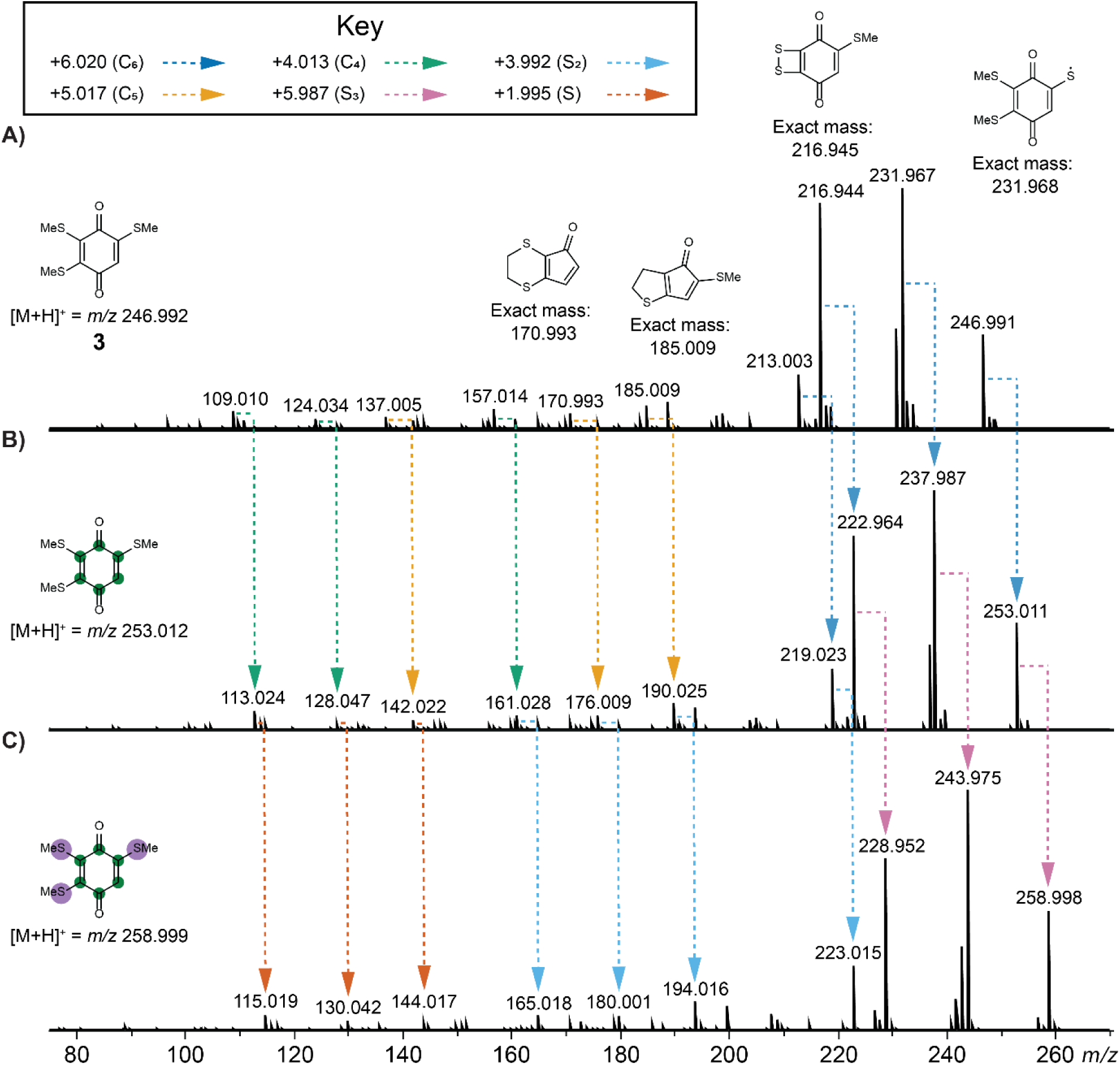
Annotated MS^2^ spectra highlighting mass shifts due to isotope labeling of T1801 B (3) when bacterial strains were cultured in M9 minimal media supplemented with A) L-methionine and L-tyrosine, B) L-methionine and L-tyrosine-^13^C_9_,^15^N, or C) L-methionine-^34^S and L-tyrosine- ^13^C_9_,^15^N.

**Figure S15.**
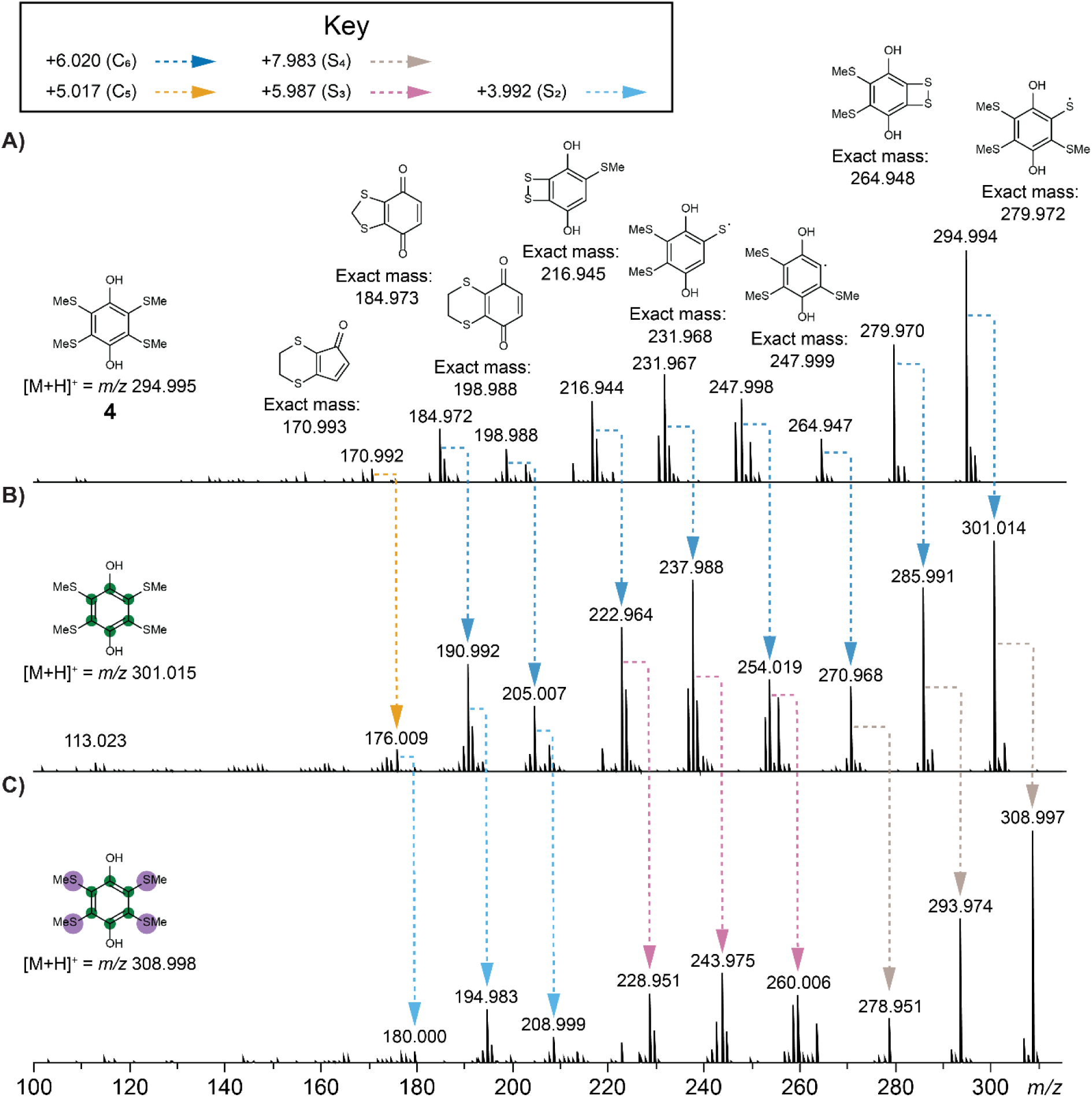
Annotated MS^2^ spectra highlighting mass shifts due to isotope labeling of T1801 C (4) when bacterial strains were cultured in M9 minimal media supplemented with A) L-methionine and L-tyrosine, B) L-methionine and L-tyrosine-^13^C_9_,^15^N, or C) L-methionine-^34^S and L-tyrosine- ^13^C_9_,^15^N.

**Figure S16.**
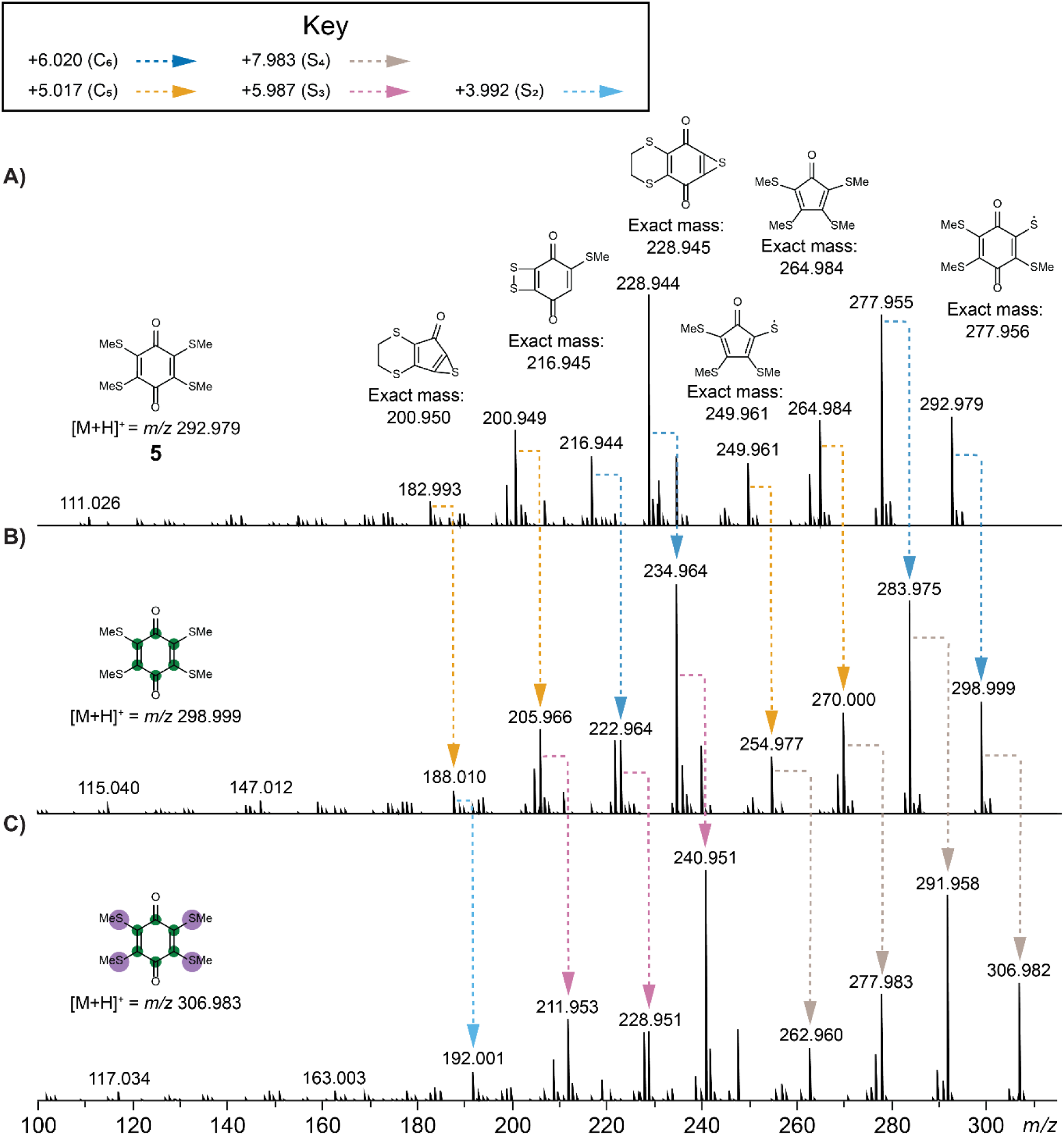
Annotated MS^2^ spectra highlighting mass shifts due to isotope labeling of T1801 D (5) when bacterial strains were cultured in M9 minimal media supplemented with A) L-methionine and L-tyrosine, B) L-methionine and L-tyrosine-^13^C_9_,^15^N, or C) L-methionine-^34^S and L-tyrosine-^13^C_9_,^15^N.

**Figure S17.**
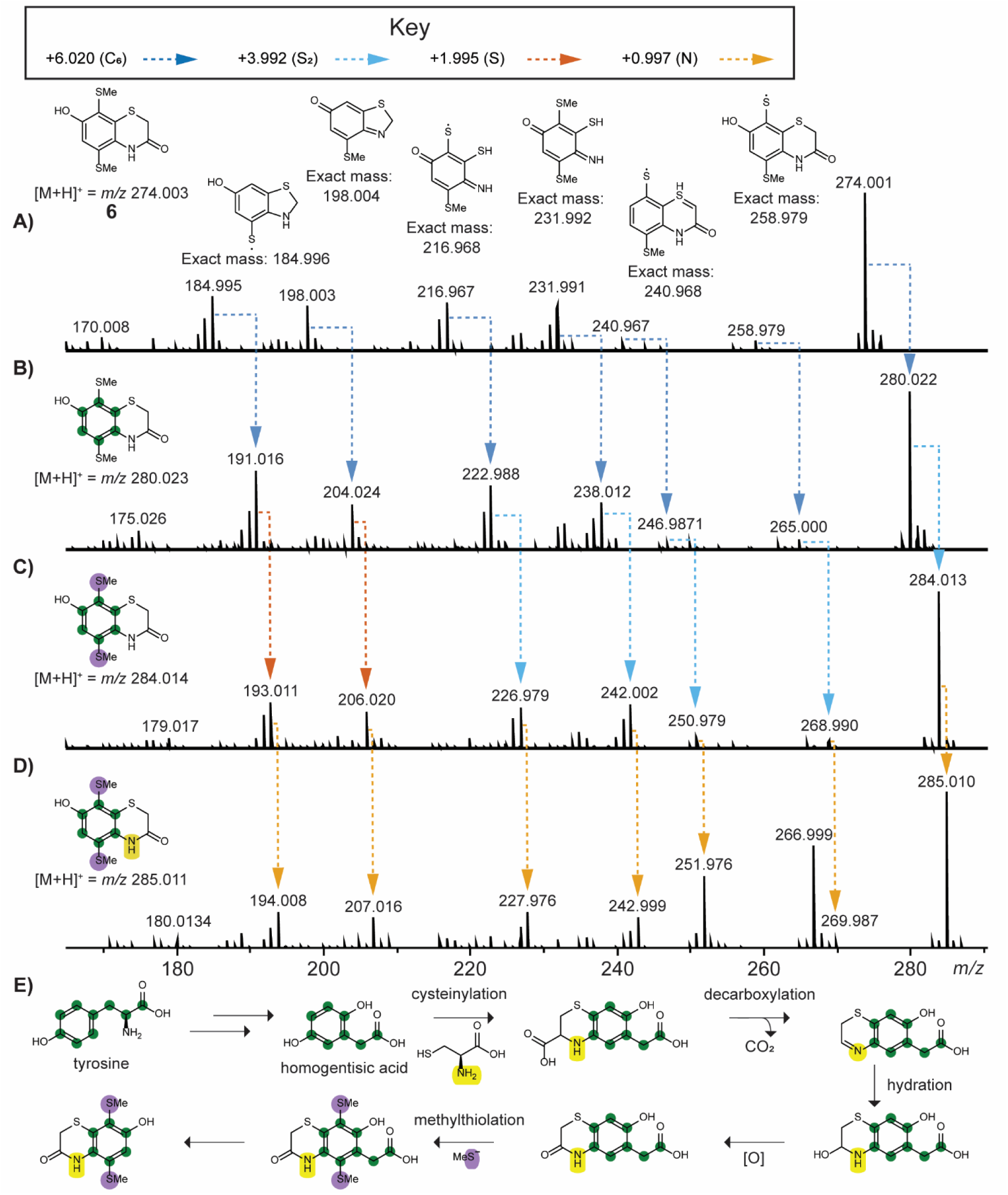
Annotated MS^2^ spectra highlighting mass shifts due to isotope labeling of BPSS2111- 2113:B (**6**) when bacterial strains were cultured in M9 minimal media supplemented with **A**) L-cysteine, L-methionine and L-tyrosine, **B**) L-cysteine, L-methionine and L-tyrosine-^13^C_9_,^15^N, **C**) L-cysteine, L-methionine-^34^S and L-tyrosine-^13^C_9_,^15^N, or **D)** L-cysteine-^15^N, L-methionine-^34^S and L-tyrosine-^13^C_9_,^15^N. **E)** Proposed putative biosynthetic scheme of **6** with tyrosine, methanethiol (via methionine), and cysteine as biosynthetic precursors supported by use of isotopically labeled amino acids. The formation of p-hydroquinone from homogentisic acid is biochemically precedented via formation of gentisate aldehyde and gentisate as intermediates. Similar pathway may be involved in the final step shown for formation of 6 and needs further investigation.

**Figure S18.**
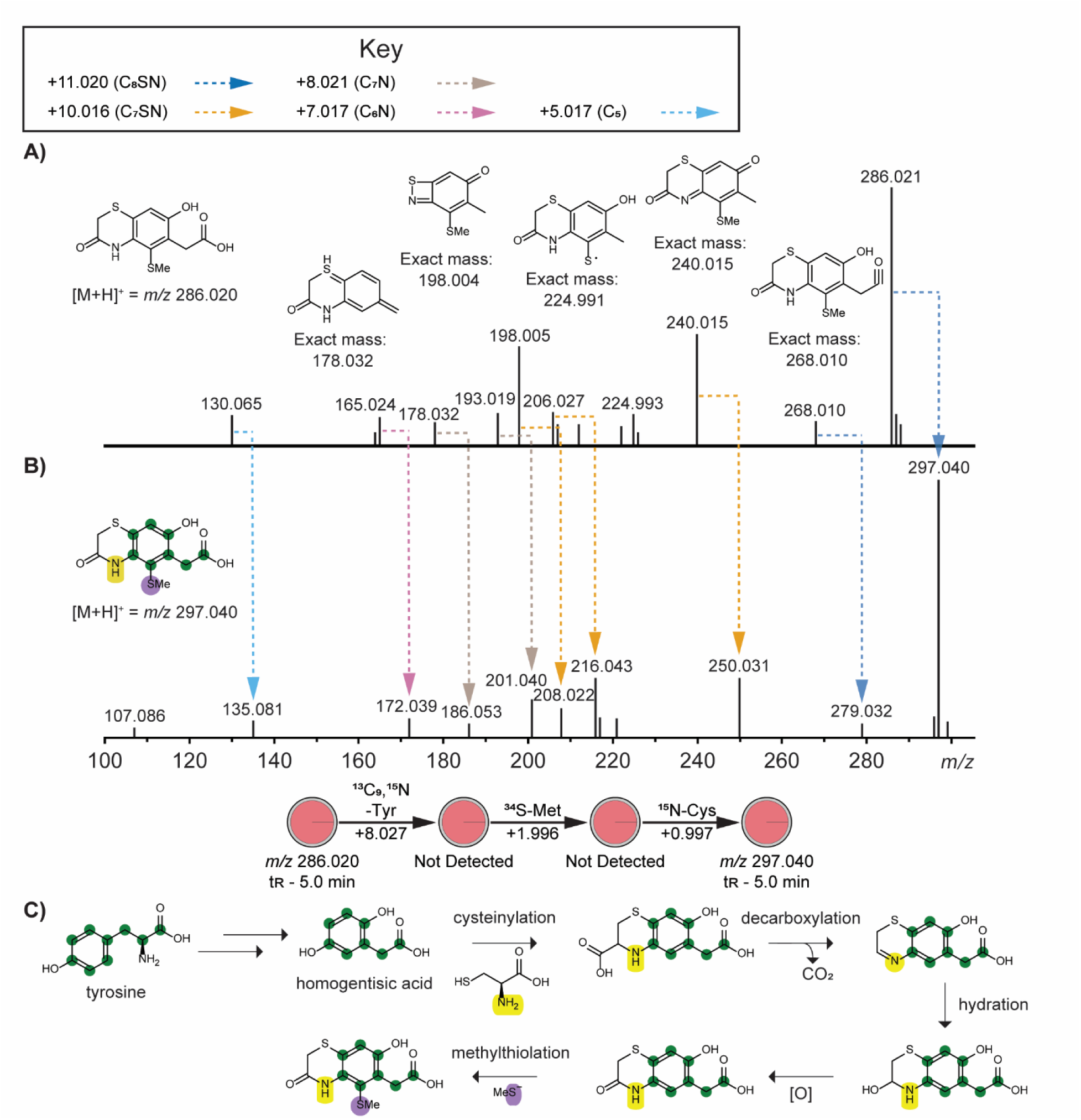
MS^2^ spectra highlighting mass shifts due to isotope labeling of *m/z* 286.020 when bacterial strains were cultured in M9 minimal media supplemented with A) L-cysteine, L- methionine and L-tyrosine or B) L-cysteine-^15^N, L-methionine-^34^S and L-tyrosine-^13^C_9_,^15^N. This metabolite was not detected without the addition of cysteine to culture media. C) Proposed putative biosynthetic scheme with tyrosine, methanethiol (via methionine), and cysteine as biosynthetic precursors supported by use isotopically labeled amino acids.

**Figure S19.**
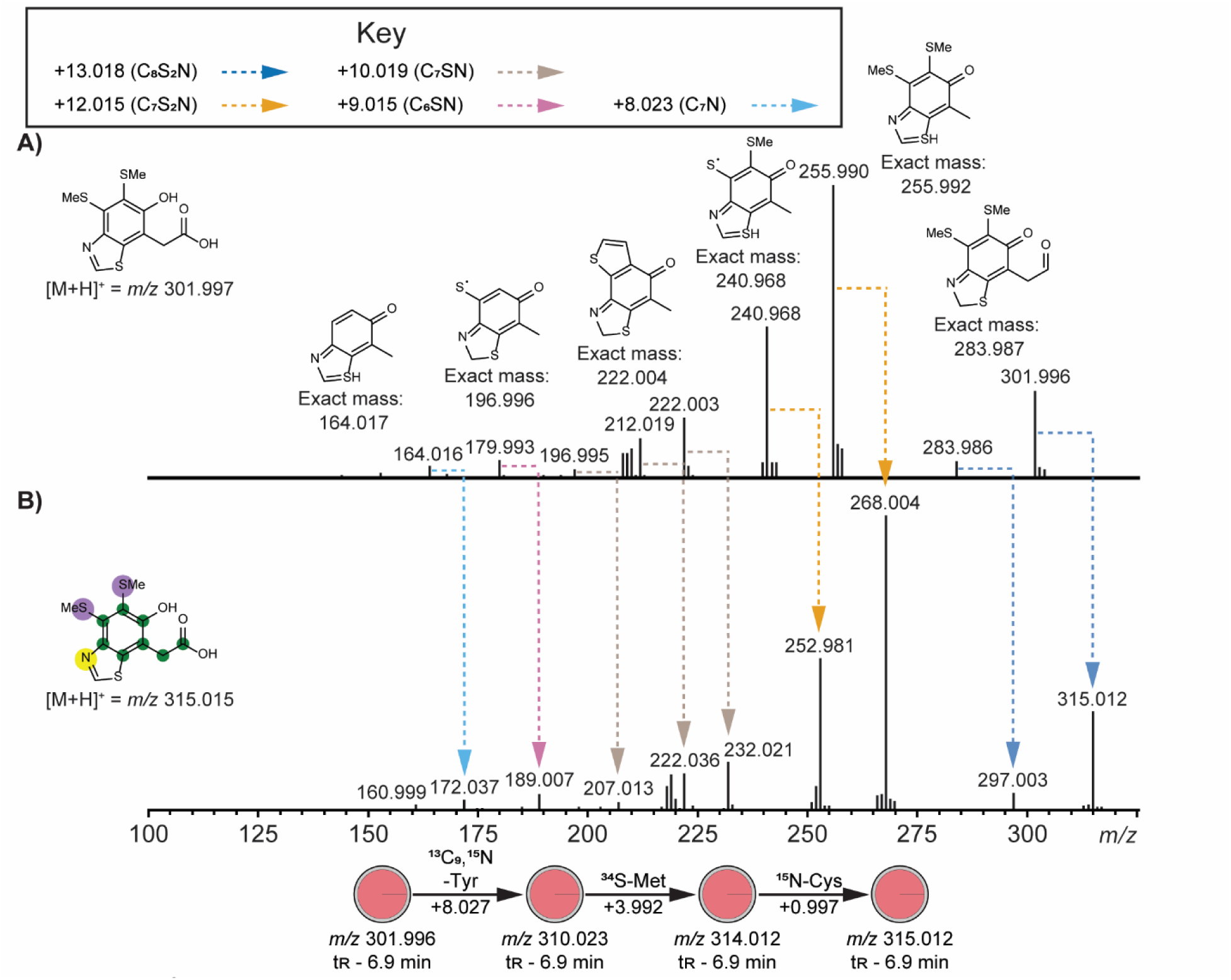
MS^2^ spectra highlighting mass shifts due to isotope labeling of *m/z* 301.997 when bacterial strains were cultured in M9 minimal media supplemented with A) L-cysteine, L- methionine and L-tyrosine or B) L-cysteine-^15^N, L-methionine-^34^S and L-tyrosine-^13^C_9_,^15^N. This compound was indeed predicted as benzothiazole by CANOPUS (Fig. 3C).

**Table S1.** Metabolite features containing the substructure represented by motif_584.

| Scan | Precursor mass | Retention time | Motif | Probability | Overlap | Annotation |
| --- | --- | --- | --- | --- | --- | --- |
| 2292 | 261.007 | 4.56 | motif_584 | 0.679608 | 0.493558 | in-source fragment, trimethylthiolated homogentisic acid with decarboxylation |
| 2251 | 289.002 | 7.45 | motif_584 | 0.675191 | 0.531873 | in-source fragment, trimethylthiolated homogentisic acid with water loss |
| 2284 | 307.013 | 7.44 | motif_584 | 0.750129 | 0.525838 | trimethylthiolated homogentisic acid |
| 1295 | 344.026 | 6.28 | motif_584 | 0.106529 | 0.439528 |  |
| 2467 | 345.036 | 6.37 | motif_584 | 0.121201 | 0.471647 |  |
| 1289 | 348.076 | 5.02 | motif_584 | 0.927003 | 0.51192 |  |
| 1762 | 349.068 | 5.12 | motif_584 | 0.258121 | 0.479237 |  |
| 1836 | 396.06 | 6.15 | motif_584 | 0.280166 | 0.502761 | Biotransformation, 261.007 + adenine |
| 2273 | 398.055 | 10.81 | motif_584 | 0.987558 | 0.514671 | Biotransformation, 261.007 + anthranilic acid |
| 2312 | 410.077 | 6.2 | motif_584 | 0.205768 | 0.457935 | Biotransformation, 261.007 + methyladenine |
| 2291 | 424.093 | 6.83 | motif_584 | 0.35599 | 0.456931 | Biotransformation, 261.007 + dimethyladenine |
| 2404 | 446.091 | 8.48 | motif_584 | 0.397729 | 0.466327 |  |
| 2299 | 464.124 | 8.81 | motif_584 | 0.321176 | 0.443457 | Biotransformation, 261.007 + isopentenyladenine |
| 2306 | 491.015 | 10.73 | motif_584 | 0.884963 | 0.486001 |  |
| 2293 | 495.122 | 5.63 | motif_584 | 0.377754 | 0.439826 |  |
| 1332 | 519.009 | 10.55 | motif_584 | 0.129938 | 0.375176 |  |
| 1348 | 551.144 | 7.96 | motif_584 | 0.643436 | 0.501734 | Biotransformation, 261.007 + trimethoprim |
| 2249 | 551.144 | 7.46 | motif_584 | 0.610395 | 0.541385 | Biotransformation, 261.007 + trimethoprim |

**Table S2.** Results of BLASTP search results, using L-methionine γ-lyase from Pseudomonas putida (UniProtKB accession: P13254) as the query sequence.

| *B. cenocepacia* J2315 |  |  |  |  |  |  |
| --- | --- | --- | --- | --- | --- | --- |
| Subject Locus Tag | Description | Max Score | Total Score | Query Cover | E-value | Percent Identity |
| BCAM0721 | O-acetylhomoserine (thiol)-lyase | 273 | 273 | 96% | 1e-88 | 40.29% |
| BCAM0999 | O-succinylhomoserine sulfhydrylase | 241 | 241 | 98% | 4e-77 | 38.6% |
| BCAL1853 | cystathionine beta-lyase | 140 | 140 | 93% | 2e-38 | 29.41% |
| BCAM0463 | cystathionine beta-lyase | 137 | 137 | 92% | 2e-37 | 30.13% |
| BCAL1502 | aspartate aminotransferase | 35.8 | 35.8 | 12% | 0.008 | 38.89% |
| *B. cenocepacia* K56-2Valvano |  |  |  |  |  |  |
| Subject Locus Tag | Description | Max Score | Total Score | Query Cover | E-value | Percent Identity |
| BURCENK562V_RS06960 | O-acetylhomoserine aminocarboxypropyltransferase | 273 | 273 | 96% | 1e-88 | 40.29% |
| BURCENK562V_RS08330 | O-succinylhomoserine sulfhydrylase | 241 | 241 | 98% | 4e-77 | 38.6% |
| BURCENK562V_RS22600 | cystathionine beta-lyase | 140 | 140 | 93% | 2e-38 | 29.41% |
| BURCENK562V_RS19900 | cystathionine beta-lyase | 137 | 137 | 92% | 2e-37 | 30.13% |
| BURCENK562V_RS24385 | aspartate aminotransferase | 35.8 | 35.8 | 12% | 0.008 | 38.89% |

